# Different combinations of laccase paralogs non-redundantly control the lignin amount and composition of specific cell types and cell wall layers in *Arabidopsis*

**DOI:** 10.1101/2022.05.04.490011

**Authors:** Leonard Blaschek, Emiko Murozuka, Delphine Ménard, Edouard Pesquet

## Abstract

Vascular plants reinforce the cell walls of the different xylem cell types with lignin, a phenolic polymer. Specific lignin chemistries are conserved between the cell wall layers of each cell type to support their functions. Yet the mechanisms controlling the tight spatial localisation of specific lignin chemistries remain unclear. Current hypotheses focus on a control by monomer biosynthesis and/or export, while their cell wall polymerisation is viewed as random and non-limiting. Here we show that cell wall polymerisation using combinations of multiple different laccases (LACs) non-redundantly and specifically control the lignin chemistry in different cell types and their distinct cell wall layers. We dissected the roles of *Arabidopsis thaliana* LAC4, 5, 10, 12 and 17 by generating quadruple and quintuple loss-of-function mutants. Different combinatory loss of these LACs lead to specific changes in lignin chemistry affecting both residue ring structures and/or aliphatic tails in specific cell types and cell wall layers. We moreover showed that the LAC-mediated lignification had distinct functions in specific cell types. Altogether, we propose that the spatial control of lignin chemistry depends on different combinations of LACs with non-redundant activities immobilised in specific cell types and cell wall layers.

## Introduction

Lignin is a complex, heterogeneous phenolic polymer that is deposited in the cell walls of specialised cell types (Meents et al. 2018). Associated with the evolutionary emergence of the plant vasculature and the transition to terrestrial habitats, lignin confers structural rigidity and hydrophobicity to the vascular system (Ménard et al. 2022; Eriksson et al. 1991). Lignin deposition proceeds in three steps: biosynthesis of monomers in the cytoplasm (Barros et al. 2015), their export into the apoplast (Perkins et al. 2019) followed by their oxidative polymerisation by radical coupling catalysed by laccases (LACs) and class III peroxidases (PRXs) in the cell wall (Blaschek and Pesquet 2021). Although defined at the singular, the term lignin describes a multitude of chemically diverse polymers. This chemical diversity comprises variation in both phenolic ring (hydroxyphenyl, **H**; guaicayl, **G**; syringyl, **S**) and aliphatic tail (alcohol, X_CHOH_; aldehyde, X_CHO_) of the canonical phenylpropanoid monomers. In addition to phenylpropanoids, other monomers incorporated into lignin include benzaldehydes (Kim et al. 2003), flavonoids (Lan et al. 2015; Rencoret et al. 2022), stilbenoids (del Río et al. 2017; Rencoret et al. 2019) as well as other residues containing phenyl (**P**) structures (Faix and Meier 1989; Kawamoto 2017). The main lignified vascular cell types in plants are tracheary elements (TEs), which act as both structural support and sap conduits for long distance water transport. Lignin is essential for structural support at the organ level, as drastic reductions lead to dwarf plants unable to stand upright (Bonawitz and Chapple 2010). Lignin is equally crucial at the cellular level, as reductions in lignin impair the capacity of TEs to withstand the negative pressures required to transport water (Ménard et al. 2022). In addition, most angiosperms and some gnetales also develop lignified xylem fibres located within (xylary fibres, XF) or between (interfascicular fibres, IF) vascular bundles, which fine-tune the mechanical properties of plant organs (Zhong and Ye 1999). To support their specific cellular functions, the concentration, composition and structure of lignins are tailored to each cell type and their different cell wall layers – chemistries that are conserved between plant species (Pesquet et al. 2019). Lignin amount is generally higher in the secondary cell walls (SCWs) of TEs than in SCWs of fibres, as well as generally higher in primary cell walls/middle lamella (CML) and cell corners (CCs) than SCWs (Serk et al. 2015). However, large variations are also observed between the different TE morphotypes, with early forming protoxylem (PX) TEs being far less lignified than later forming metaxylem (MX) TEs (Ménard et al. 2022). Lignin composition, depending on the proportion of monomers with different ring structures, varies drastically between cell types and cell wall layers, with fibre SCWs enriched in **S** residues, TE SCWs enriched in **G** residues, and CML/CCs at the interface of these two cell types enriched in **H** residues (Terashima and Fukushima 1988; Fukushima and Terashima 1990). Similarly, the proportion of monomers with different aliphatic tails is also specifically controlled between cell types and cell wall layers, as well as between morphotypes, with high levels of X_CHO_ in MXs compared to PXs, and high levels of X_CHO_ in CMLs compared to SCWs (Peng and Westermark 1997; Blaschek, Champagne et al. 2020). The lignification of each cell type moreover progresses by incorporating distinct lignin residues at specific phases of the development and maturation of each cell type, exemplified by **S** and X_CHO_ residues that are mostly incorporated in the late maturation phases of TEs compared to **H** and G_CHOH_ (Terashima and Fukushima 1988; Kutscha and Gray 1972; Blaschek, Champagne et al. 2020). The temporal control of cell wall lignification is specific to each cell type. TEs lignify their SCWs mostly after having committed programmed cell death (Pesquet et al. 2010; Pesquet et al. 2013; Smith et al. 2013) whereas fibres gradually and centripetally lignify their SCWs as they are being deposited (Terashima et al. 1996; Fukushima and Terashima 1991; Barros et al. 2015). The mechanisms enabling such a specific spatio-temporal control of lignin chemistry in different cell types and cell wall layers are still unclear.

The current hypotheses for the mechanisms explaining this cell type/cell wall layer specific control of lignification mainly rely on the regulation of monomer biosynthesis and/or their export to cell walls, while their polymerisation is mostly viewed as random, non-limiting and unspecific. Our current view of lignin formation thus considers that the oxidation of lignin monomers and their subsequent cross-coupling is only guided by thermodynamics and steric hindrance (Blaschek and Pesquet 2021). However, once in the apoplast, lignin monomers are extremely mobile due to autocrine, paracrine (Smith et al. 2017; Smith et al. 2013) and potentially endocrine secretions (Aoki et al. 2019; Blaschek, Champagne et al. 2020). This cell–cell transport is essential for TEs, which lignify *post-mortem* (Ménard et al. 2022; Pesquet et al. 2010; Pesquet et al. 2013; Derbyshire et al. 2015) by polymerising the monomers secreted by adjacent living cells. Neighbouring fibres also use these externally supplied monomers semi-cooperatively in complement to their own export during and after the deposition of SCW polysaccharides (Barros et al. 2015; Blaschek, Champagne et al. 2020). The high mobility of lignin monomers in the cell wall contrasts with the spatially restricted differences of lignin chemistry in each cell wall layer of each cell type. This dissonance suggests that additional mechanisms control lignification at the level of the cell wall itself. Such spatial control could depend on different LACs and/or PRXs with distinct catalytic activities that are targeted and immobilised in specific cell types and cell wall layers (Derbyshire et al. 2015; Chou et al. 2018). In fact, different LACs show highly distinct binding pocket and active site topologies, suggesting differences in catalytic efficiency, pH optimum and/or substrate specificity (Blaschek and Pesquet 2021). Indeed, LACs involved in the formation of specialised lignins in seed coats and compression wood were recently shown to exhibit a certain degree of substrate specificity for monomers with different ring substitutions (Wang et al. 2020; Hiraide et al. 2021; Zhuo et al. 2022). The role of LAC specificity in the main developmental lignification of xylem however remains unclear. Herein, we provide experimental evidence demonstrating that the lignin polymerisation capacity of the different cell wall layers of each xylem cell type spatially controls its lignin chemistry. Using higher-order loss-of-function mutants of *LACs* involved in vascular lignification, we show that the isolated activity of different *LAC* paralogs is specific to each cell type and cell wall layer, and discriminately incorporates specific monomers. The resulting differences in lignin chemistry also affected its function in distinct cell types, distinctly affecting the cell wall mechanical resistance of TEs to negative pressure and the hydrophobicity of fibre cell walls. Altogether, we show that different immobilised combinations of LAC paralogs with specific activities control the lignin chemistry in each cell wall layer and cell type to support their different functions.

## Materials & methods

### Plant material

*Arabidopsis thaliana* plants were grown in three independent growth instances, consisting of 5 plants per genotype each, between January 2020 and July 2021 from stratified (2 d at 4°C) seeds in 1:3 vermiculite:soil in E-41L2 growth cabinets (Percival, USA). Growth conditions were cycled in a 1:6:1:16 h dusk:night:dawn:day program. Relative humidity was kept at 50%, irradiation were set to 50, 0, 50, 100 μmol m^−2^ s^−1^, and temperature at 20, 18, 20, 22°C for dusk, night, dawn and day respectively. The following insertional single mutants in the Col-0 background were crossed to obtain higher-order mutants: *lac4-2* (GK-720G02-025278; Berthet et al. 2011), *lac5-1* (SALK_063466; Cai et al. 2006), *lac10-1* (SALK_017722; Cai et al. 2006), *lac11-1* (SALK_063746; Zhao et al. 2013), *lac12-2* (SALK_125379), *lac17-1* (SALK_016748; Berthet et al. 2011). Plants were confirmed to be homozygous mutants by genomic DNA extraction from two green rosette leaves using the EZNA Plant DNA kit (Omega Bio-Tek, D3485) followed by PCR genotyping using standard *Taq* DNA polymerase (18038-042, Invitrogen) to amplify sequences unique to WT and mutant alleles using specific primer combinations (table S1). PCRs used 10 ng of genomic DNA per 10 μl reaction and were cycled through 98°C (3 min), [98°C (45 s), 57°C (45 s), 72°C (90 s)] ×35, 72°C (5 min). Amplicons were separated using agarose gel electrophoresis with mini Gel II (VWR), stained with Midori Green (MG 04, Nippon Genetics) and imaged using UV-transillumator (Biorad). Descendants of plant crosses were planted and genotyped until the desired homozygous mutant combination was obtained. Three independent crosses to obtain the double homozygous mutants *lac10-1×lac11-1* and *lac12-2×lac11-1* were unsuccessful. Homozygous phenylpropanoid mutants used as references in figure 4 were *fah1* (EMS mutant; Meyer et al. 1998), *ccr1-3* (SALK_123-689; Mir Derikvand et al. 2008) and *cad4;5* (Blaschek, Champagne et al. 2020). After 9 w of growth, plants were harvested. The basal 2 cm of the primary inflorescence stem were stored in 70% ethanol at −20°C until sectioning. The stem bases were vacuum infiltrated with ultrapure water, embedded in 10% agarose (A9539, Sigma-Aldrich), and sectioned to 50 μm thickness using a VT1000s vibratome (Leica). Sections were then stored in 70% ethanol at −20°C and rehydrated when needed with ultrapure water. Seeds for the root growth assay were surface sterilised using 70% ethanol followed by 5% bleach (Klorox), plated on on 0.8% agar-agar (20767.298, VWR) containing 0.5× MS medium (pH 5.7) including vitamins (M0222, Duchefa Biochem), and stratified for two days in darkness at 4°C. Seedlings were grown for 11 d in 60% relative humidity and 150 μmol m^−2^ s^−1^ in 16:8 h day:night conditions at 18 and 24°C respectively.

### Growth phenotyping

Growth kinetics of *A. thaliana* was analysed by image-based phenotyping. Images were acquired every 2 to 3 days under even illumination in front of a black background with a Nikon D750 camera equipped with a Sigma 50mm F1.4 DG HSM lens. An X-Rite ColorChecker Classic Mini colour card was included in each image as a colour and size reference. Images were segmented and analysed using a custom script within the PlantCV (v. 3.8) framework (Gehan et al. 2017). Used scripts are available at https://github.com/leonardblaschek/plantcv. To account for slight differences in the timing of inflorescence stem bolting between growth instances, the stem growth kinetics were aligned at the time of bolting (approximated as the day the plant surpasses a height of 5 cm). Additional manual phenotyping was done at the time of harvest after 9 w of growth, measuring stem number, primary stem height, primary stem width at base, and silique length for 5 mature siliques from the primary stem.

### Laccase activity assays

Activity assays in sections were performed by vacuum infiltrating 50 μm transverse stem cross sections (stored in 70% ethanol at −20°C) with water for 2 hours and placing them in 0.1 mM sodium acetate (pH 4, 5) or sodium phosphate (pH 6, 7) buffer in a MatTek no. 1.5 glass-bottom 96-well plate. Sections for negative controls were previously autoclaved at 121°C for 15 min using a Tuttnauer 3870EL autoclave. The final reaction volume was 200 μl and substrate concentrations were 0.07 mM DAF, 0.5 mM DAB or 5 mM PYGL (table S2). Real-time images were taken every 10 (DAF, PYGL) or 15 (DAB) min using a Zeiss Axiovert 200M equipped with a 25× objective (NA 0.4), an Axiocam 506 color camera and an automated stage and shutters. Each section was imaged in three Z-positions (spaced approximately 10 μm apart) to account for any drift in the Z-axis. Images were aligned in Fiji (Schindelin et al. 2012) using the SIFT plugin, transformed into absorbance and measured in 20 circular areas (0.7 μm diameter) per image and cell type. Measurements were background corrected by subtracting the absorbance of the unlignified phloem. To compare the activities across different pH, the non-enzymatic substrate oxidation rates in autoclaved sections were subtracted from the oxidation rates measured in the untreated sections in figures 2A and S3D. Comparison of activities at similar pH (figure 2B,C) did not require any subtraction.

**Figure 1.**
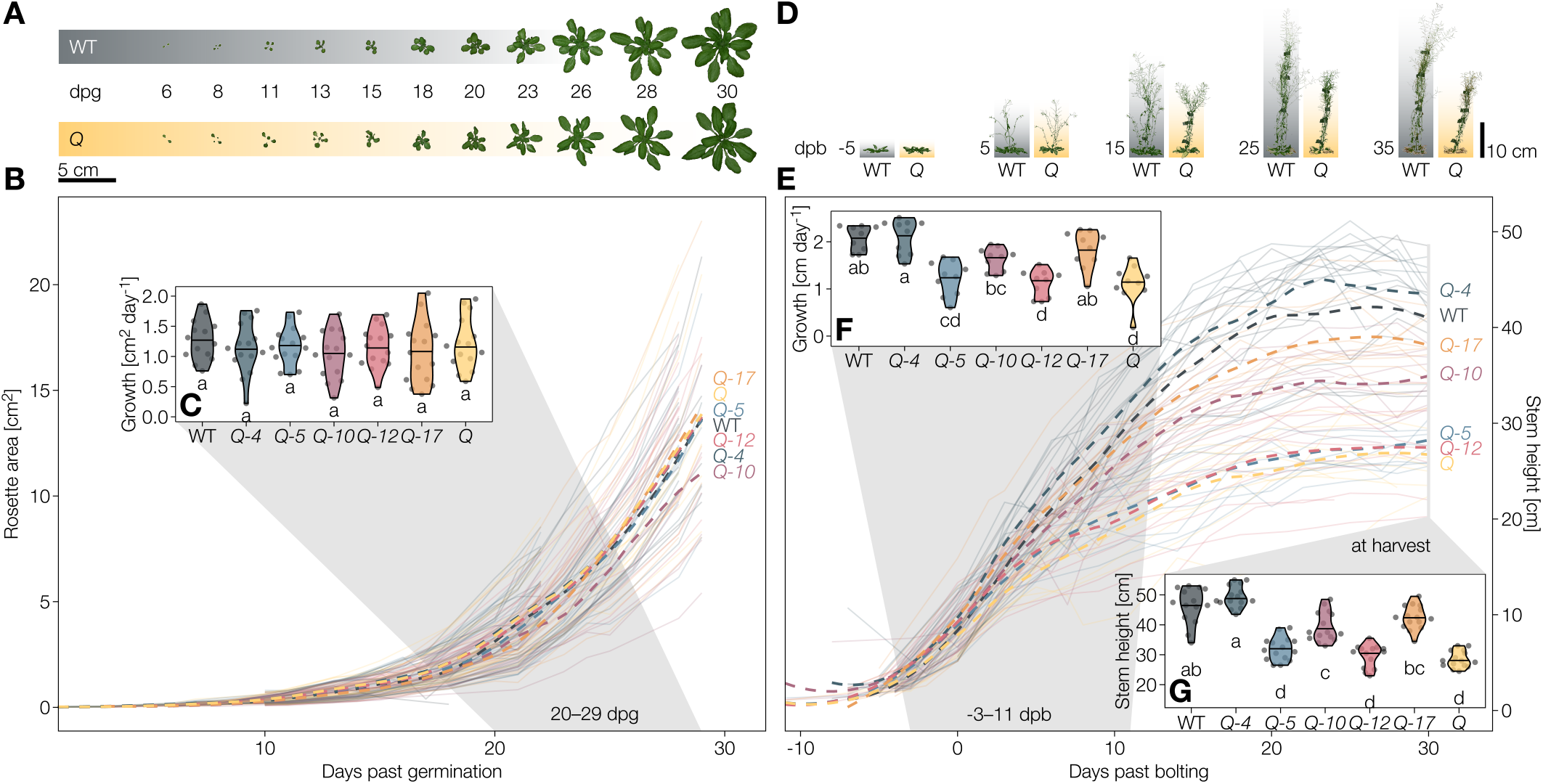
Phenotypical characterisation of higher-order *lac* mutants. **A** Representative rosettes of the WT and the *Q* mutant between six days past germination (dpg) and bolting at around 30 dpg. **B** Projected area of the rosettes from germination to bolting. **C** Growth rates of the projected rosette areas between 20 and 29 dpg. **D** Representative inflorescence stems of the WT and the *Q* mutant from 5 days before bolting to 35 days past bolting (dpb). **E** Projected stem height from bolting to senescence. **F** Growth rates of the projected stem height from −3 to 11 dpb (defined as reaching a height of > 5 cm). **G** Stem height (stretched length of the primary inflorescence stem) at harvest. Solid lines represent individual plants, dashed lines are moving regressions (LOESS) showing the genotype averages. Different letters indicate statistically significant differences between genotypes according to a Tukey-HSD test (*α* = 0.05); *n* = 10–15 individual plants from 2–3 independent growth instances per genotype.

**Figure 2.**
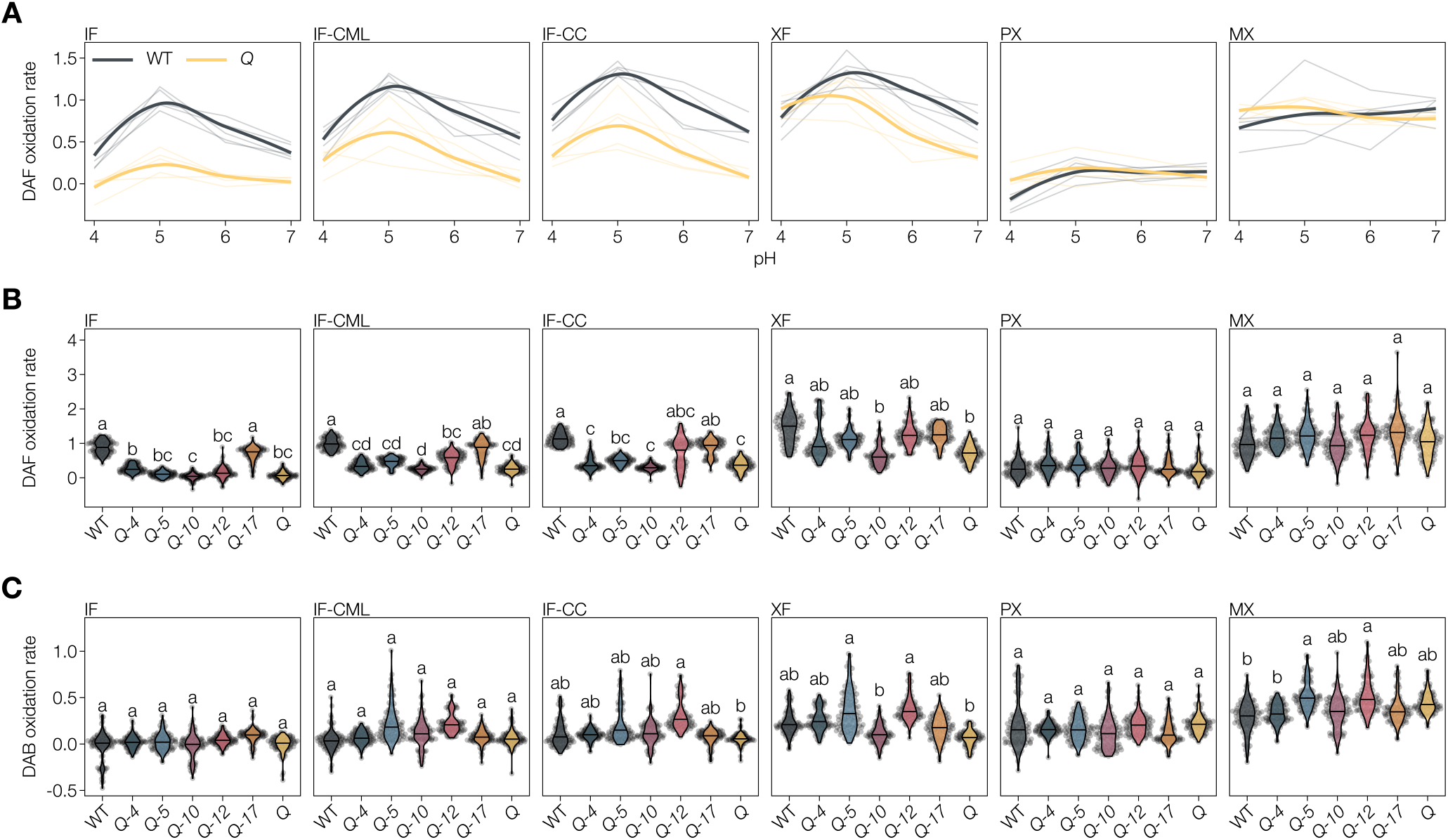
Laccase activity in the cell walls of higher-order laccase mutants. **A** Activity of WT and *Q* towards DAF at pH 4–7, expressed as relative substrate oxidation rates. Thin lines represent 5 individual replicates, thick lines are the overall average by local regression (LOESS). Oxidation rates in autoclaved sections are subtracted to show only LAC mediated oxidation. **B** Activity of WT and higher-order *lac* mutants towards DAF, expressed as relative substrate oxidation rates. **C** Activity of WT and higher-order *lac* mutants towards DAB, expressed as relative substrate oxidation rates. Different letters indicate statistically significant differences between genotypes according to a Tukey-HSD test (per panel; *α* = 0.05); 20 measurements were done for each cell type in each of *n* = 5 individual plants per genotype.

The corrected absorbance values were plotted against time to identify the linear phase of activity following the initial unspecific staining and before the plateauing of absorbance values at the end of the time course. From the linear phase, substrate oxidation rates were determined by the slope of each linear regression.

### Histochemical analysis

#### Wiesner stain

Cross-sections of the stem bases were stained with the Wiesner test as described previously (Blaschek, Champagne et al. 2020). Briefly, hydrated cross-sections were imaged in water using a Zeiss Axiovert 200M, equipped with a EC Plan-Neofluar 40x objective (NA 0.75) and an automated stage, then stained by removing the cover slip and dropping 40 μl of 0.5% phloroglucinol (P3502, Sigma-Aldrich) in 1:1 95% ethanol:37% HCl onto the section, and imaged again after 2 min incubation. Images were corrected for uneven illumination using a picture of an empty slide and stitched in Fiji (Preibisch et al. 2009). Absorbance values were obtained by aligning unstained and stained images, transforming the images into uncalibrated optical density and measuring circular areas (0.45 μm diameter) of cell walls or specific cell wall layers for each cell type. The specific absorbance of the stain was obtained by subtracting the unstained absorbance from the stained absorbance for each measured point. Lastly, the absorbance measurements were corrected for unspecific tissue clearing by the ethanolic acid in the Wiesner reagent and/or changes in illumination by subtracting the absorbance difference between stained and unstained images of the unlignified phloem from all measurements.

#### Mäule stain

Hydrated cross-sections of the stem bases were incubated in 0.75% KMnO_4_ (223468, Sigma-Aldrich) in water for 5 min, rinsed twice with water, incubated in 3M HCl for 1 min, rinsed twice with water, and finally incubated in 10% NaCO_3_ (71360, Sigma-Aldrich) for roughly 30 s. The sections were mounted in 10% NaCO_3_ between glass slide and coverslip and imaged within 10 minutes after staining. Imaging and stitching were performed as described for the Wiesner stain. The hue of the stain was estimated by transforming the RGB image into the HSB colourspace and measuring circular areas (0.45 μm diameter) of cell walls or specific layers in the hue channel. The resulting values were divided by 255 and multiplied by 360 to transform them from the 8-bit range to the 360° hue scale.

### Raman microspectroscopy

Raman microspectroscopy was performed on hydrated crosssections in ultrapure water between glass slide and cover slip, using a LabRAM HR 800 (Horiba, France) equipped with a Nd:YAG 532 nm laser and a 50x objective (NA 0.42). Spectra were acquired with a spectral resolution of 2 cm^−1^ from 100 to 1800 cm^−1^. Acquisition parameters were kept constant, using a grating of 600 grooves mm^−1^, a laser power of 56 mW and 100 accumulations of 3 s per measurement. Asymmetric-least-squares baseline correction was performed as previously described by Blaschek, Nuoendagula et al. (2020). Cellulose crystallinity was estimated according to (Agarwal et al. 2010). For the non-ratiometric lignin compositional analyses, spectra were normalised to the total lignin signal. To ease the interpretation of the Raman band intensities and ratios, measurements from well described phenylpropanoid mutants were included: the **S**-free *fah1* mutant (EMS; Meyer et al. 1998), the lignin deficient *ccr1-3* mutant (SALK_123-689; Mir Derikvand et al. 2008) and the **G**_CHO_ overaccumulating *cad4;5* double mutant (Blaschek, Champagne et al. 2020).

### Lignin autofluorescence

Lignin autofluorescence in the IF was measured in hydrated cross-sections mounted between glass slide and coverslip in 50% glycerol. Acquisition was performed using Zeiss LSM780/800 confocal laser scanning microscopes equipped with a Plan-Apochromat 20× M27 objective (NA 0.8), a 405 nm diode laser and long-pass emission detection >410 nm according to Decou et al. (2017). For direct comparison of fluorescence intensities (figure S6B), imaging parameters were kept constant between genotypes. For the relative quantification of cell wall layer-specific autofluorescence (figure 5C,D) imaging parameters were adjusted between genotypes to optimise signal to noise ratio.

**Figure 3.**
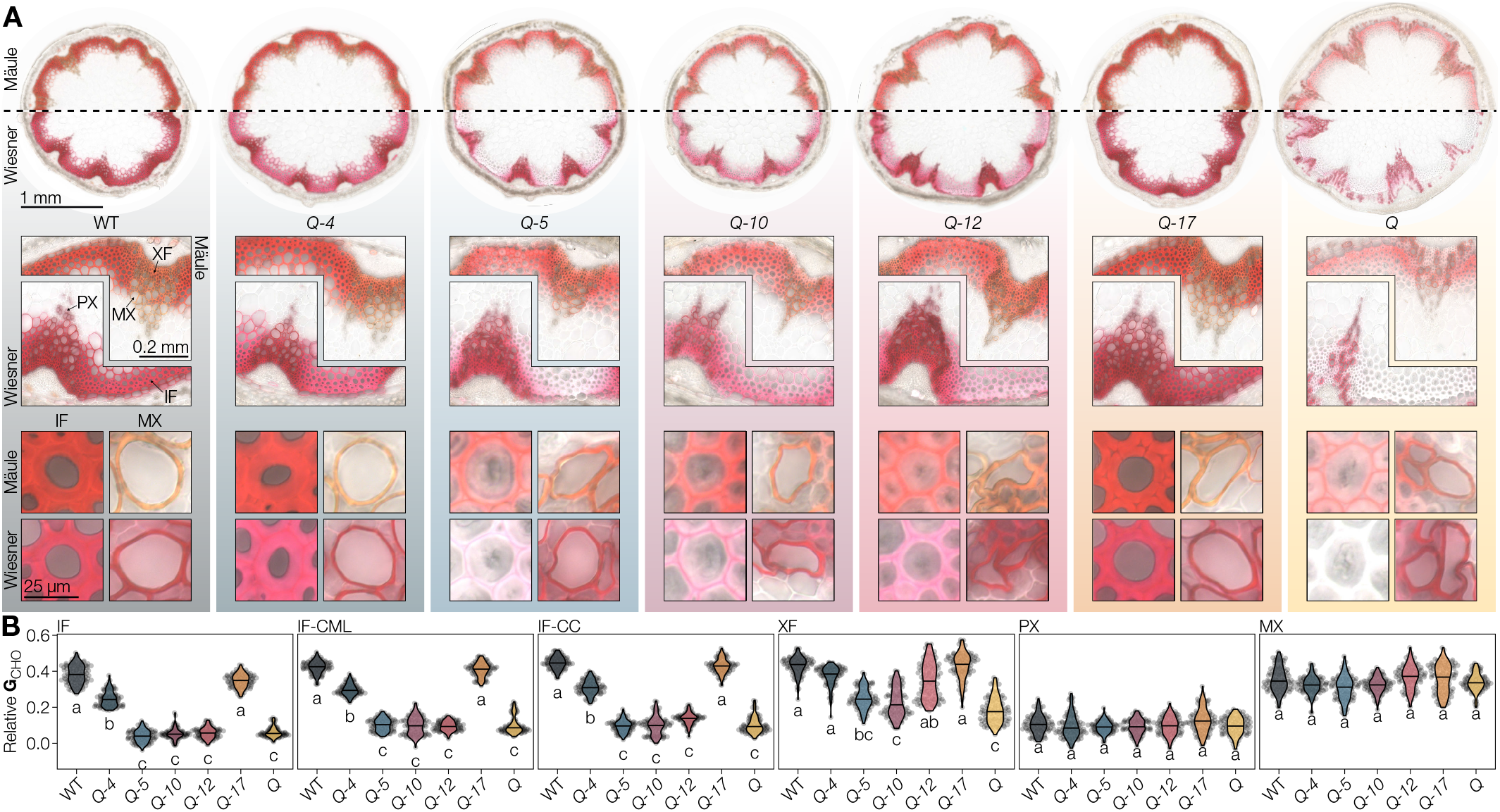
Histology of higher-order *lac* mutants. **A** Mäule and Wiesner stained transverse stem sections at increasing levels of detail: full sections, vascular bundles with adjacent interfascicular fibres (IF), and single cells of IF and metaxylem (MX). **B** Relative coniferaldehyde concentrations in the IF, the compound middle lamella of the IF (IF-CML), xylary fibres (XF), protoxylem (PX), and MX. Different letters indicate statistically significant differences between genotypes according to a Tukey-HSD test (per panel; *α* = 0.05); 20 measurements were done for each cell type in each of *n* = 5 individual plants per genotype from 2 independent growth instances.

**Figure 4.**
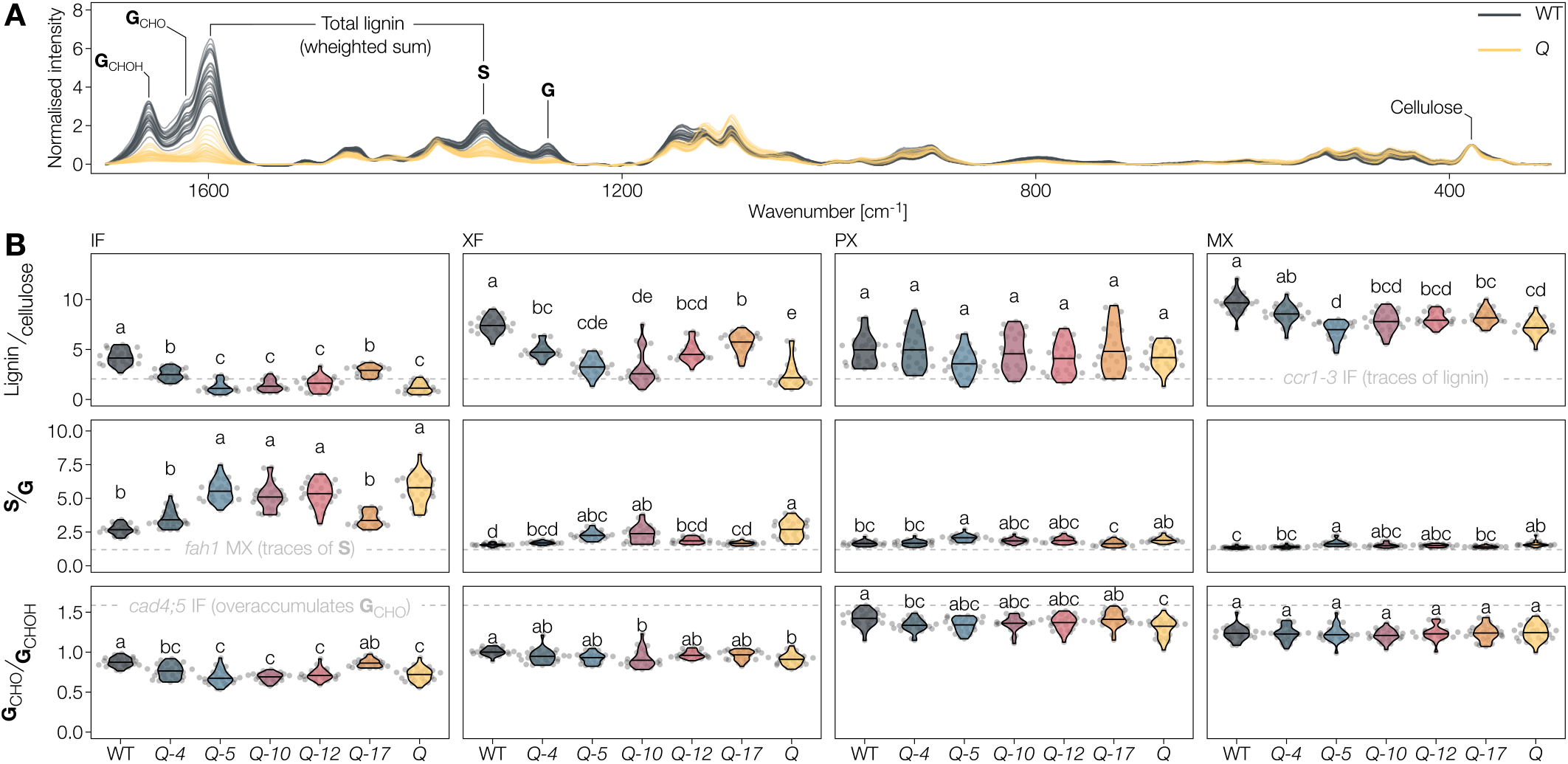
Lignin composition in higher-order *lac* mutants measured by Raman microspectroscopy. **A** Raman microspectra from the IF of the WT and *Q* mutant with bands used for quantification. Spectra are scaled to the cellulose band at 378 cm^−1^. **B** Ratiometric characterisation of lignin amount and composition. Because of the undefined Raman scattering coefficients of the different lignin substituents, the ratios are relative, not absolute. References from well characterised phenylpropanoid loss-of-function mutants are indicated by dashed grey lines to ease interpretation. Different letters indicate statistically significant differences between genotypes according to a Tukey-HSD test (per panel; *α* = 0.05); spectra of 5 individual cells were measured for each cell type in each of *n* = 5 individual plants per genotype from 2 independent growth instances.

**Figure 5.**
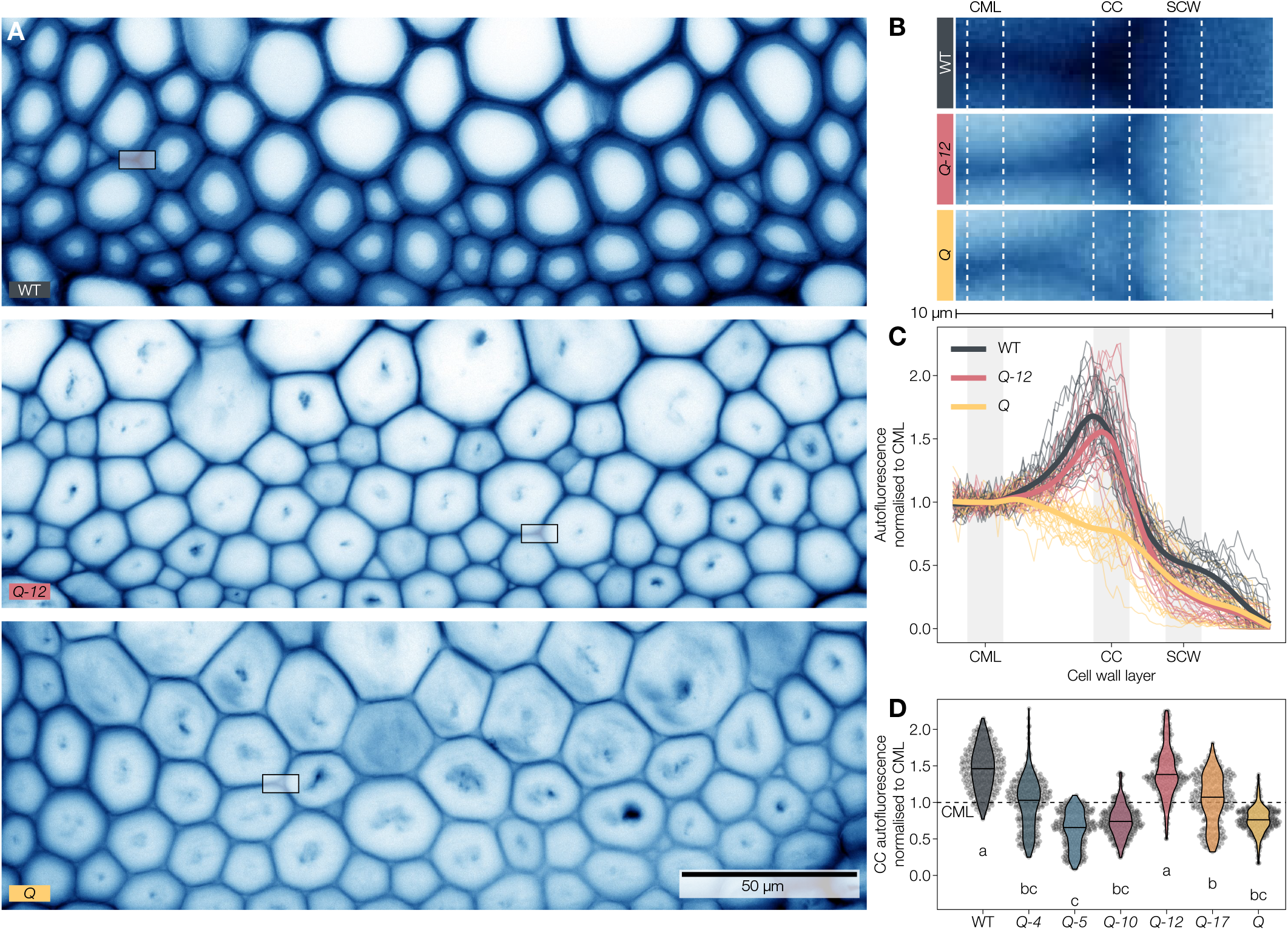
Lignin autofluorescence in the cell corners (CCs) depends on LAC12. **A** Lignin autofluorescence in the IF of the WT, *Q-12* and *Q*. Contrast of the images is adjusted for visibility. **B** Magnification of the boxes drawn in panel A, showing the compound middle lamella, CC and SCW of two adjacent IFs. Contrast of the images is adjusted for visibility. **C** Line profiles of autofluorescence intensity normalised to the compound middle lamella. Thin lines are individual profiles from 5 independent plants, the thick line is an average by local regression (LOESS). **D** Normalised fluorescence intensity measured in the shaded stretches indicated in panels B and C. Different letters indicate statistically significant differences between genotypes according to a Tukey-HSD test (per panel; *α* = 0.05); each point represents one pixel from a total of 25 line profiles in *n* = 5 individual plants per genotype from 2 independent growth instances.

### TE collapse

The collapse of each TE morphotype was measured from the images of Wiesner stained sections (see above). TE perimeters were traced in ImageJ (Schindelin et al. 2012) using the “better wand tool” macro (https://gist.github.com/mutterer/4d3c831ca6a7e698f77eba0261b086c5) when the image contrast allowed it, otherwise by freehand selection. Pixel coordinates of the perimeters were loaded into R (v. 4.10) and kernel-smoothed using the package “smoothr” (v. 0.2.2) to avoid shape artefacts where the perimeter followed the image pixelation. Area and convex hull of the smoothed perimeters were calculated using the package “sf” (v. 1.0-6), convexity represents the ratio of area to convex hull.

### Cell wall swelling

Hydrated cross-sections of the stem bases were incubated in 0.01% Astra blue (B0770, Sigma-Aldrich) in water to stain cellulose, rinsed in water, incubated in 0.01% Safranin-O (1.15948, Merck) in water to stain lignin, rinsed in water, and finally mounted between glass slide and coverslip in water to simultaneously stain all cell walls (Srebotnik and Messner 1994). Imaging and stitching were performed as described for the Wiesner stain. After imaging, the stained sections were let to air dry at room temperature overnight and subsequently imaged again. Finally, the sections were rehydrated by incubating them in water in the fridge for several days, mounted between glass slide and coverslip in water, and imaged again. Images of each section in the three stages were aligned in ImageJ. The thickness of 10 cell walls per section was measured in each state, at a point equidistant from the adjacent cell corners.

### Data analysis and visualisation

If not described otherwise, all data analysis and visualisation was done in R (v. 4.10), relying on the “tidyverse” (v. 1.3.1) collection of packages (Wickham et al. 2019). Tukey-HSD and Kruskal-Wallis tests were performed using the average values of each biological replicate (individual plants, specified in each figure legend as “*n* = *x*”) using the “tukey-grps” package (https://github.com/leonardblaschek/tukeygrps), which uses functions from the “stats” and “dunn.test” packages. The R code to reproduce all analysis and figures is available at https://github.com/leonardblaschek/Rscripts/blob/master/2021_lac_paper.rmd.

## Results

### LAC paralogs non-redundantly and synergistically affect stem growth

To discriminate which of the 17 laccase paralogs are implicated in the lignification of secondary cell walls in xylem cells, we analysed available gene expression data (micro-array and RNA sequencing) from *A. thaliana* inducible pluripotent cell cultures triggered to differentiate into TEs (Derbyshire et al. 2015). Candidate *LAC* paralogs were selected based on high co-expression levels with SCW formation marker gene *CesA7/IRX3* and TE autolysis marker gene *XCP2* (Ménard et al. 2015). This approach identified *A. thaliana LAC*s *4, 5, 10, 11, 12* and *17* as co-up-regulated during TE SCW formation and potentially implicated in vascular lignification (figure S1). Previous promoter:reporter studies confirmed the expression of all six identified candidates in cambium and/or xylem (Turlapati et al. 2011). The large number of implicated *LAC*s likely explains why previous studies using single loss-of-function mutants did not detect any obvious phenotypic changes (Cai et al. 2006), except for a slight *irregular xylem* (*irx*) phenotype in *lac4/irx12* (Brown et al. 2005; Berthet et al. 2011). Indeed, significant decreases in plant height were only observed in plants that had lost at least three *LAC* paralogs (figure S2A). To expose the functions of each individual LAC candidate, we generated homozygous quintuple (*Q*) loss-of-function mutant of LACs *4, 5, 10, 12* and *17*. We excluded *LAC11* because its loss in triple mutants together with *LAC4* and *17* results in sterile plants (Zhao et al. 2013). Additionally, we were unable to obtain homozygous double mutants of *LAC11* in combination with either *LAC10* or *LAC12.* To define the specific functions of each individual LAC, we analysed the quadruple loss-of-function mutants *lac5;10;12;17* (*Q-4*), *lac4;10;12;17* (*Q-5*), *lac4;5;12;17* (*Q-10*), *lac4;5;10;17* (*Q-12*) and *lac4;5;10;12* (*Q-17*) in comparison to the *Q* mutant and wild-type (WT) plants. Primary root growth in seedlings and their overall morphology were unaffected by the stacking of *lac* mutations (figure S2B). For soil-grown plants, we performed kinetic analyses using image-based phenotyping of the growth of rosette leaves and inflorescence stems (figure 1). The growth of rosette leaves and the timing of bolting were unaltered by stacking *lac* mutations (figure 1A–C, S2C). In contrast, stem growth rate, final height, and silique length of *Q* were reduced to roughly 50%, 60% and 90% of the WT respectively (figure 1D–G, S1D). The reduced growth rate and final height were restored to WT levels completely in *Q-4* and *Q-17*, and partially in *Q-10* but remained uncompensated in *Q-5* and *Q-12* (figure 1E–G). Silique length was fully restored to WT levels in Q-4, *Q-10* and *Q-17*, partly restored in *Q-5* but not compensated in *Q-12* (figure S2D). Our results indicated a major and non-redundant intervention of *LAC4, LAC10* and *LAC17* in both stem and silique development in contrast to *LAC5* and *LAC12*. Overall, higher-order *lac* mutations mainly altered the vertical growth of inflorescence stems that suggested roles in structural support and long-distance water transport.

### LAC activity is cell type- and substrate-specific

To understand how the genetic modulation of *LACs* resulted in the different mutant phenotypes observed, we measured LAC activity in higher-order *lac* mutants. LAC activity has been previously performed *in situ* but without any temporal resolution or quantitative read-out (Hiraide et al. 2021). We used real-time imaging to measure oxidation rates directly in the cell walls of different cell types in ethanol-cleared and rehydrated stem cross-sections (figure S3). This real-time *in situ* activity assay also enabled us to evaluate substrate-specificity, using a set of synthetic compounds with either phenylamine or phenolic groups shown to be oxidised by LACs (Hiraide et al. 2021; Richardson et al. 2000; Bao et al. 1993), as well as define optimal pH. Autoclaved sections served as negative controls and showed very little non-enzymatic activity, except for the increased auto-oxidation of PYGL at higher pH (figure S3). With the phenylamine 2,7-diaminofluorene (DAF), LAC activity peaked at pH 5 in most cell types in WT except for MX TEs which exhibited a stable LAC activity from pH 4–7, whereas PX showed almost no LAC activity (figure 2A). The magnitude of activity at the different pH was reduced in the *Q* mutant for SCW, CML and CC of IFs, slightly for XFs but unaltered for MX TEs (figure 2A). The observed pH optimum however depended on substrate, as using the phenolic pyrogallol (PYGL) showed two activity optima at pH 5 and 7, which differed between cell wall layers and cell types (figure S3). These results showed that cell walls of each cell type differed in their LAC substrate oxidation capacity and optimal pH. Activity analyses using DAF – which was more reliable in detecting differences between genotypes than PYGL – at optimal pH for higher-order mutants revealed that each LAC affected specifically the activity profiles of the different vascular cell types (figure 2B). In the different cell wall layers of IFs, LAC activity towards DAF was almost abolished in the *Q* mutant and only consistently restored in *Q-17* (figure 2B). Additionally, *Q-12* showed increased levels of LAC activity specifically in CCs and to a lesser extent in the CML (figure 2B). LAC activity towards DAF in both PX and MX TEs remained unchanged in all mutants (figure 2B). In contrast, XFs exhibited an intermediate LAC activity for DAF between IFs and TEs, with a significant reduction in the *Q* mutant, restored partly by the different LACs except LAC10 (figure 2B). Additional analyses to evaluate differences in substrate specificity of the different LACs were made using a second phenylamine substrate, 3,3-diaminobenzidine (DAB). Activity towards DAB in the cell wall layers of IFs showed little differences between genotypes except for specifically increased CC activity in *Q-12* (Figure 2B). MX TEs showed increased activity in *Q-5* and *Q-12* (Figure 2B), showing that LAC5 and 12 were active towards DAB in contrast to the other LAC paralogs. Lastly, XF also presented enhanced activity in *Q-5* and *Q-12* similarly to MX TEs but also a reduced activity in *Q-10* and *Q* compared to WT plants (figure 2B). Our results thus confirmed the effectiveness of each insertional mutation by specifically reducing LAC activity. Altogether, the different higher-order *lac* mutants revealed the specific localisation in distinct cell wall layer and cell types of LAC paralogs. Despite their functional overlap, LAC paralogs were only partially redundant in terms of oxidation rate, substrate range, optimal pH condition and spatial localisation in cell walls and cell types.

### Each LAC paralog has cell type-specific effects on lignification

To evaluate how the observed cell type and cell wall layer-specific differences in LAC activity affected lignification, we performed histochemical *in situ* analyses of lignin. Staining extractive-free stem cross-sections of 9 w old plants using either the Mäule or the Wiesner tests enabled cell wall layer level resolutions directly in whole plant organ biopsies (figure 3A). The Mäule test detects changes in the proportions of different ring structures by staining **S** residues in red and other lignin constituents in brown (Yamashita et al. 2016). Mäule stained WT plants showed the expected intense red in the **S**-enriched fibre SCWs and brown in the essentially **S**-depleted TEs (figure 3A). higher-order *lac* mutants showed weak red staining in the SCWs of IFs of *Q-5, Q-10, Q-12* and *Q* in contrast to *Q-4* and *Q-17* (figure 3A), indicating LAC4 and LAC17 are the predominant enzymes ensuring **S** residue accumulation in IF SCWs. However, CMLs of IFs remained strongly red stained in all genotypes (figure 3A and S4), indicating a more robust lignification mechanism in the primary wall than the SCW layers. Compared to the WT, slight red-shifts of 5–10 hue degrees were apparent for all mutants except *Q-17* in IFs and for *Q-5, Q-10, Q-12* and *Q* in TEs, indicating increases in **S**/**G** ratio (figure S4). The Wiesner test specifically detects coniferaldehyde (**G**_CHO_) and can be used to reliably quantify **G**_CHO_ concentration *in situ* (Blaschek, Champagne et al. 2020). IF SCWs, CMLs and CCs showed almost no quantifiable stain in *Q-5, Q-10, Q-12* and Q, as well as a reduction in *Q-4* when compared to WT and *Q-17* (figures 3A, B). This result revealed the predominant role of LAC17, with a minor contribution of LAC4, on the incorporation of **G**_CHO_ in the SCWs of IFs. In contrast, MX TEs appeared similarly stained to the WT plants for all genotypes (figure 3B), revealing that LACs are implicated in regulating certain residues differently between cell types. XFs were affected similarly but not as severely as IFs by higher-order *lac* mutation, showing largely reduced but not abolished accumulations of **G**_CHO_ in *Q-5, Q-10* and *Q* in contrast to *Q-4, Q-12* and *Q-17* (figure 3B). This observation indicated that LAC4, LAC17 and LAC12 could restore **G**_CHO_ accumulation in XFs to WT levels. Although PX TEs had much lower **G**_CHO_ levels than MX TEs in WTs, they remained similarly unaffected in all mutants (figure 3B). Even though we could not monitor changes in total lignin amounts using these methods, our histochemical analyses showed that specific LAC paralog combinations are responsible for distinctively controlling the incorporation of lignin residues with different ring structures and/or aliphatic tails between cell wall layers and cell types.

### Each LAC paralog non-redundantly determines lignin composition in the different cell types

To quantify the specific effects of each *LAC*s on both lignin amount and composition in the different cell types, we used Raman microspectroscopy on extractive-free cross-sections of higher-order *lac* mutants. Several Raman bands can be used for the relative quantification of different cell wall polymers including total lignin and cellulose amounts, lignin **S**/**G** ratio and total **G**_CHOH_ to terminal **G**_CHO_ ratio (figure 4A) (Blaschek, Nuoendagula et al. 2020; Yamamoto et al. 2020). As our Raman set-up did not have the spatial resolution to distinguish between SCW and CML, we observed in WT plants the highest concentration of lignin in MX TEs, followed by XFs and then IFs and PXs (figure S5A). In WT plants, lignin **S**/**G** ratio was highest in IFs, followed by XFs and PXs, and then MXs (figure S5A), whereas lignin **G**_CHO_/**G**_CHOH_ ratio was highest in PXs, then MXs, followed by XFs and then IFs (figure S5A).

Lignin to cellulose amount was reduced for all lignified cell types in all genotypes compared to WT, except for PX that remained unaltered (figure 4B). The broad changes in lignin in the *Q* mutant also affected the structure of other cell wall polymers, reducing cellulose crystallinity in XFs (figure S5B). Among the quadruple mutants, *Q-4* and *Q-17* had the highest lignin concentrations in all tested cell types. Slight differences between these two mutants suggested a more prominent role in MX lignification for LAC4 and fibre lignification for LAC17. *Q-5, Q-10* and *Q-12* revealed smaller, but specific roles of the respective LACs depending on the cell type. IF SCWs of *Q-5, Q-10, Q-12* and *Q* had lignin to cellulose ratios lower than the phenylpropanoid mutants *ccr1-3* (Mir Derikvand et al. 2008), showing that IF lignin concentration was highly dependent on LAC4 and LAC17 (figure 4B). In XFs, *Q-5* and *Q-10* restored lignin accumulation less than *Q-12* which reached *Q-4* and *Q-17* levels (figure 4B), indicating an important redundant contribution of LAC12, LAC4 and LAC17 compared to the minor contribution of LAC5 and LAC10 to XF lignin accumulation. In MXs, reductions were less pronounced and showed no differences between *Q-5* and Q, whereas *Q-10* and *Q-12* showed a slight increase (figure 4B). Altogether, our results show clearly that different LAC combinations, varying in paralogs and relative contribution, control lignin concentration in the SCWs of the different cell types of plant vascular tissues.

Lignin **S**/**G** ratio was increased in all cell types of the *Q* mutant compared to WT plants, and the different higher order *lac* mutants exhibited intermediate profiles varying for each LAC in magnitude and cell type (figure 4B). SCWs of all cell types had **S**/**G** ratios greater than the phenylpropanoid mutant *fah1-2* which only has traces of **S** residues (Meyer et al. 1998). These results confirmed our histochemical analysis using the Mäule test (figure 3A). The **S**/**G** changes were likely due to a large reduction in **G** residues rather than an increase of **S** residues (figure S5B). In fibres, **S**/**G** increases in IFs were restored to WT levels in *Q-4* and *Q-17*, whereas **S**/**G** increases in XFs were restored to WT levels by *Q-4, Q-17* and *Q-12* (figure 4B). In sap conducting TEs, the **S**/**G** increases in PXs and MXs were restored to WT levels in *Q-17* but also *Q-4, Q-10* and *Q-12* but not Q-5, revealing here a similar contribution of LACs for controlling lignin ring structure composition. As the **S**/**G** ratio differs greatly between TEs and fibres, our results indicate that different LAC paralog combinations control the ring structure proportions in these two different categories of cell types.

The ratio of terminal **G**_CHO_ to total **G**_CHOH_ groups was reduced in all cell types of the *Q* mutant compared to WT, except for MX (figure 4B) but did not reach the **G**_CHO_ over-accumulation levels of the phenylpropanoid mutants *cad4;5* (Blaschek, Champagne et al. 2020; Yamamoto et al. 2020). These results confirmed our histochemical analysis using the Wiesner test (figure 3B). Intermediate profiles were obtained differing for each higher-order mutant in its magnitude and cell type. In fibres, **G**_CHO_/**G**_CHOH_ decreases in IFs were restored to WT levels fully by *Q-17* and partly by *Q-4* and not in other genotypes, whereas **G**_CHO_/**G**_CHOH_ decreases in XFs were partly restored in *Q-4, Q-5, Q-12* and *Q-17* but not *Q-10* (figure 4B). In sap conducting TEs, **G**_CHO_/**G**_CHOH_ decreases in PXs were fully restored to WT levels by *Q-17*, partly by *Q-5, Q-10* and *Q-12* and slightly by Q-4, whereas MXs were not affected (figure 4B). The fact that the ratio of terminal **G**_CHO_ to **G**_CHOH_ was not restored by LAC4 in IFs or PXs corroborates its reduced propensity to incorporate **G**_CHO_ compared **G**_CHOH_ to shown by the Wiesner stain (figures 3B, 4B). Together, these results show that different combinations of LACs control the accumulation of residues with specific aliphatic functions in different cell types.

higher-order *lac* mutants also affected the incorporation of non-canonical residues, such as benzaldehydes and residues with phenyl (**P**) rings, differently between cell types. The proportion of **P** residues was increased in *Q* mutants for both fibre types but not in TE morphotypes (figure S5B). These increases were differently compensated by the different LAC paralogs, with **P** residues fully restored to WT levels in IFs of *Q-4* and *Q-17* and in XFs of Q-4, Q-5, *Q-12* and *Q-17* (figure S5B). Benzaldehyde residues were also increased in *Q* but only in IFs, which could be restored to WT levels by all LAC paralogs except LAC5 (figure S5B). These results indicated that specific LAC paralogs also altered the incorporation of non-canonical residues in specific cell type. Altogether, our results clearly showed that different LACs are active in distinct cell types and cell wall layer, where they act in combination to control lignin chemistry.

### Distinct LAC paralogs are responsible for cell-wall-layer specific lignification

To further investigate the cell wall layer-specific roles of different LAC paralogs indicated by the activity assays and histochemical tests (figures 2, 3), we quantified cell wall layer-specific lignin accumulation in higher-order *lac* mutants using the UV-excited autofluorescence of lignin combined with confocal microscopy (Decou et al. 2017). This technique had a greater spatial resolution than our Raman analysis set-up that did not enable to clearly distinguish between cell wall layers. As expected, lignin autofluorescence in WT IFs was highest in CCs, intermediate in the CML and lowest in S1/S2 and S3 layers of SCWs (figures 5A–C and S6). Mirroring the results of histochemical and Raman spectroscopy analyses (figures 3, 4), SCW autofluorescence was drastically reduced in *Q-5, Q-10, Q-12* and *Q*, whereas *Q-4* and *Q-17* exhibited similar fluorescence levels as WT plants (figure S6). Spatial analyses of SCWs to distinguish S1/S2 and S3 layers revealed that *Q-4* had WT-like lignin content in the S3, but not the S1/S2 layers, unlike *Q-17* that restored lignin concentration to WT levels across all layers of the SCWs (figure S6). In the primary cell wall layers, CML autofluorescence was drastically reduced in *Q-5, Q-10, Q-12* and *Q* but only slightly reduced in *Q-4* and similar to the WT in *Q-17* (figure S6). In contrast, the autofluorescence of CCs in both *Q-4* and *Q-17* was reduced to levels similar to the WT CML, while *Q-12* showed higher autofluorescence than *Q* and *Q-5*, and *Q-10* showing intermediate levels (figure S6). We then measured line profiles through the different cell wall layers to compare changes of lignin autofluorescence across the cell wall layers of single neighbouring cells (figures 5B–D and S6). To control for biological and technical variation in absolute fluorescence between plants and cells, we scaled each line profile to the average of its CML fluorescence. This approach confirmed that LAC12 activity was restricted to CCs and responsible for the specific increase in CC lignin autofluorescence levels relative to the CML (figure 5C, D). In *Q-5, Q-10* and Q, lignin autofluorescence in CCs was even lower than in CMLs (figure 5D), suggesting that the LACs and/or PRXs responsible for the residual lignification of these cell wall layers were active in CMLs but not CCs. These results showed that the lignification of IF cell wall layers depended on LAC4 and LAC17 for the S3 layer of their SCWs, LAC17 together with a minor contribution of LAC4 for the S1/S2 and CML, and LAC4, LAC12 and LAC17 for CCs. LAC12 was by itself capable of fully restore the ratio of CC to CML lignin autofluorescence to the level observed in WT plants (figures 5D and S6; Serk et al. 2015). Altogether, and in addition to the cell type and substrate specificity shown above between LAC paralogs, our results indicate that specific combinations of LAC paralogs non-redundantly control the spatially stratified lignification between CCs, CML, S1/S2 and S3 layers in vascular cells.

### Specific LAC paralogs control lignification to ensure the mechanical resistance of sap conducting cells

Lignin is essential for the proper vascular function of sap conducting cells by conferring the mechanical reinforcement required to sustain the negative pressure associated with sap transport (Ménard et al. 2022). We investigated whether the mechanical resistance of TEs was controlled by specific LACs by measuring the extent of their inward collapse in higher-order *lac* mutants. Convexity – a shape descriptor specifically characterising inwards collapse and not just general deformation (Ménard et al. 2022) – was measured for the different TE morphotypes in stem cross-sections. TE collapse varied between genotypes as well as between TE morphotypes, revealing again that different LAC paralogs had non-redundant contribution in controlling lignin-dependent mechanical resistance of specific TEs (figure 6). The early forming PX TEs showed some occasional collapse in all quadruple mutants but were only significantly collapsed in the absence of all five tested *LAC*s (figure 6). This result indicated that all investigated LAC paralogs synergistically contributed to the mechanical reinforcement of PXs. The later forming MX TEs collapsed severely in *Q*, *Q-5* and *Q-12*, to lesser extent in *Q-10* and *Q-17*, and almost not at all in *Q-4* compared to the WT (figure 6). This showed that lignin-dependent mechanical reinforcement of the MX heavily depended on LAC4, with intermediate contributions of LAC17 and LAC10, a minor role of LAC5 but no contribution of LAC12. Lastly, in the secondary xylem TEs (SX), no statistically significant TE inward collapse was observed in any of our *lac* mutants (figure S7). This confirmed previous results showing that the SX was generally less prone to inward collapse (Ménard et al. 2022), and suggested that the mechanical strengthening of SX cell walls was less dependent on the tested LACs than in PX and MX. Overall, these results show that LAC paralogs non-redundantly contribute to TE lignin-dependent mechanical reinforcement, differing in both their level of contribution and TE morphotypes, with LAC4 playing the most predominant role.

**Figure 6.**
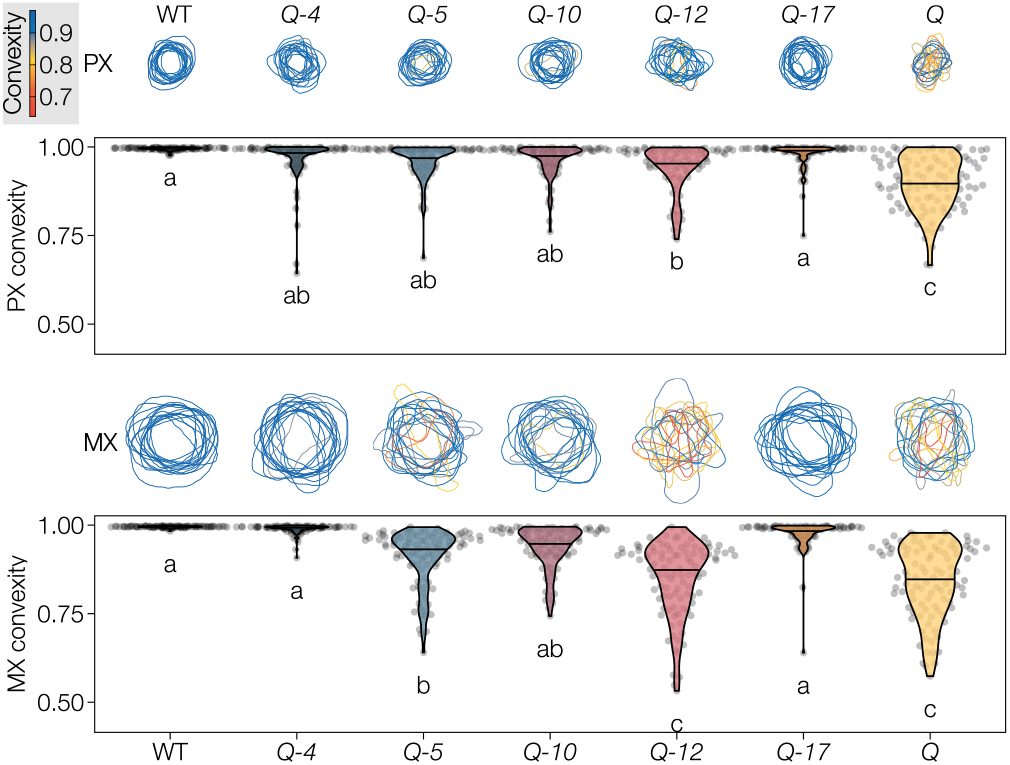
TE perimeters and their degree of inwards collapse (expressed as convexity) in the different higher-order *lac* mutants. Twenty TEs of each cell type in each of *n* = 5 individual plants per genotype were measured. The drawn outlines represent fifteen randomly sampled TEs per cell type. Different letters indicate statistically significant differences between genotypes according to a Tukey-HSD test (per panel; *α* = 0.05).

### Specific LAC paralogs control lignification to control fibre swelling capacity

The stiffness and flexibility of angiosperm plant stems is controlled by lignified TEs as well as lignified fibres. Although the stiffness of TE cell walls was significantly impacted by LAC mutations, both TE perimeter and their cell wall thickness, independently of their hydration level, were not altered by any of the mutations (figure S8). In contrast, IF SCWs in *Q, Q-5, Q-10* and *Q-12*, all with reduced lignin amounts, were significantly thicker than the more lignified IF SCWs in WT, *Q-4* and *Q-17* plants (figure 7). To test whether this increase in cell wall thickness was due to the deposition of additional cell wall polysaccharides or a lignin dependent change in cell wall properties, we stained sections with Safranin-O/Astra blue and imaged them fresh, after drying, and after rehydrating (figure 7). This allowed us to assess the proportion of IF cell wall thickness dependent on hydration. The reduced image contrast in the dried state unfortunately prevented a similar assessment of the small, densely packed XF cells. The SCWs in dried samples of all mutants presented the same thickness as WT IFs (figure 7). This result indicated that lignin controlled the hydration level of SCWs, which regulated their thicknesses (figure 7). Rehydrating the sections confirmed this observation as SCWs of IFs in *Q, Q-5, Q-10* and *Q-12* swelled up to almost the same thickness as before drying (figure 7). WT, *Q-4* and *Q-17* however had unchanged IF cell wall thicknesses in all three conditions. These results show that lignification in fibres of vascular tissues acted as a waterproofing “varnish” to stabilise the thickness of cell walls by controlling their hydration levels, with predominant and redundant roles of *LAC4* and *LAC17*.

**Figure 7.**
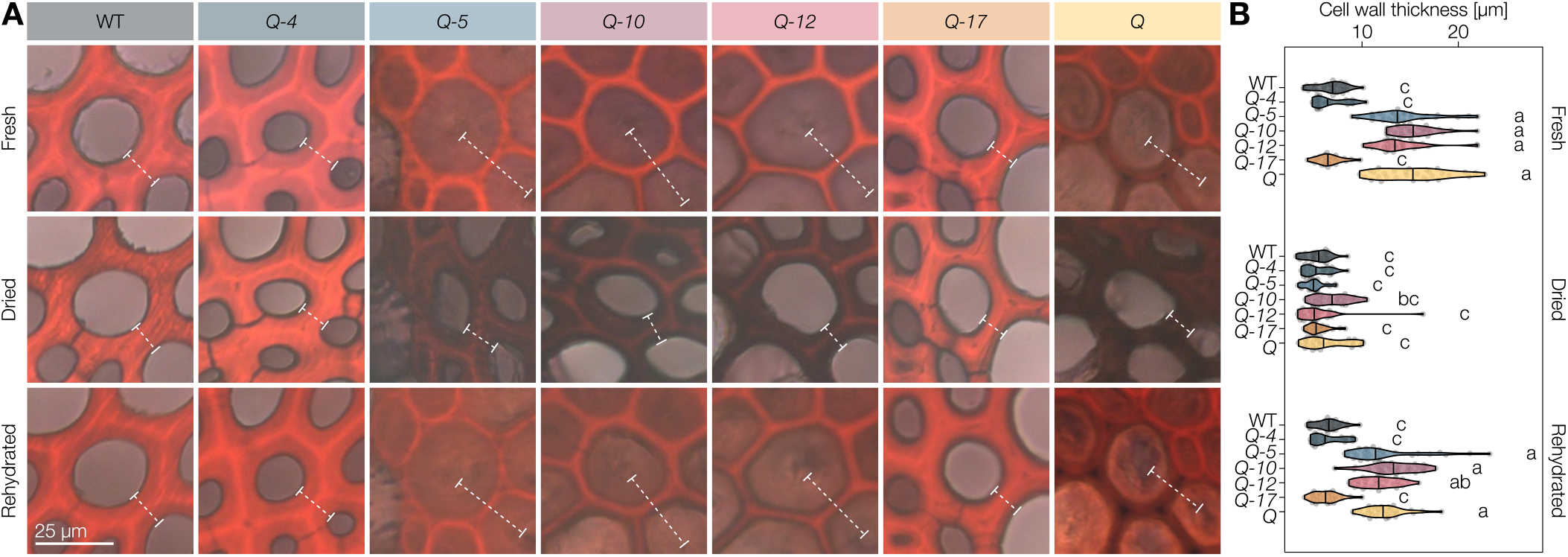
Lignin depleted IF secondary cell walls in higher-order *lac* mutants swell in water. **A** Representative Astra Blue–Safranin stained IF cell walls imaged fresh (never dried), dried (after air drying at room temperature over night) and rehydrated (after incubating the dried sections in water). The thickness of the cell wall (lumen to lumen) is indicated by dashed lines. **B** The thickness of 10 IF cell walls (lumen to lumen) from each of *n* = 3 individual plants per genotype in the three states. Different letters indicate statistically significant differences between genotypes and states according to a Tukey-HSD test (per panel; *α* = 0.05).

## Discussion

Our study highlights the necessity to measure lignin at the cellular and subcellular levels to avoid the averaging errors made when analysing whole organs. We were able to effectively show both concentration and compositional differences of lignin between cell wall layers and cell types, indicating the importance and diversity of spatial distribution of lignin between cells in plant organs. An even more precise understanding of lignification can be generated with approaches distinguishing the temporal changes occurring for each cell types during their development/maturation, such as measured using real-time imaging of inducible pluripotent cell cultures (Pesquet et al. 2010; Derbyshire et al. 2015) and/or cells along wood growth rings (Blaschek, Champagne et al. 2020; Sundell et al. 2017) and/or different stem segments from the apical meristems (Hoffmann et al. 2020; Ménard et al. 2022; Morel et al. 2022). The development of *in situ* lignin quantification methods necessary for this type of analysis is relatively recent and has yet to overcome substantial hurdles. Recent efforts have focused on indirect methods for *in situ* lignin analyses using synthetic compound feeding (Tobimatsu et al. 2013; Lion et al. 2017; Zhu et al. 2021; Morel et al. 2022). Additionally, multiple stains interacting more or less specifically with lignin have been widely used (Bond et al. 2008; Pesquet et al. 2005; Kapp et al. 2015; Donaldson and Williams 2018). However, the mechanisms controlling the incorporation of fed compounds as well as the nature and specificity of the lignin substructures interacting with specific dyes are not understood. Fluorescent labelling of lignin with safranin in *lac4;17* cross-sections showed increases in fluorescent labelling of MX TEs, revealing changes in lignin composition (Baldacci-Cresp et al. 2020) which were explained by our analysis. Ectopic transdifferentiation of PX TEs from epidermal cells in *lac4;17* mutants was previously used to demonstrate the capacity of LAC4 and 17 to incorporate fluorescent-**G**_CHOH_ (Schuetz et al. 2014). However, these feeding experiments could not define the effect on native lignin content in PX TEs of the mutant, unlike our Raman analysis (figure 4). The direct *in situ* quantification methods used in this work also have drawbacks. The label-free Raman microspectroscopy used herein had limited spatial resolution and could not reliably detect **H**, **S**_CHO_ or **S**_CHOH_ canonical residues (Blaschek, Nuoendagula et al. 2020). The direct Wiesner test, while relying on a well-defined chemical reaction (Blaschek, Champagne et al. 2020), can only detect **G**_CHO_ and is distorted by out-of-focus light absorption and the birefringence of weakly stained cell walls, restricting its detection limit and spatial resolution within cell wall layers. We additionally extended the technical possibilities of these *in situ* analyses to include time-resolved LAC activity measurements directly in cell walls of extractive-free plant cross-sections, which showed both differential activity rates, distinct substrate preference and different pH optima profiles between cell types and cell wall layers (figure 2). Single cell analyses like the *in situ* quantitative imaging methods used herein are crucial to characterise the cellular heterogeneity of lignin that gets lost through averaging in whole organ analysis and represent a necessary development to reliably understand lignin formation at the subcellular level during the development of each cell type.

The precise spatio-temporal control of lignification is pivotal for normal development (Zhao et al. 2013), drought resistance (Lima et al. 2018; Ménard et al. 2022), and defence against herbivores and pathogens (Joo et al. 2021; Whitehill et al. 2016). The importance of lignin monomer biosynthesis in each cell type (Barros et al. 2019; Blaschek, Nuoendagula et al. 2020) and their transport from the cytosol into the apoplast (Perkins et al. 2019; Väisänen et al. 2020) are essential to control lignin but insufficient to regulate the strict spatial distribution of lignin in between cell types and cell wall layers.

In addition to their extreme mobility in cell walls, alcohol and aldehyde phenylpropanoids have been shown to diffuse freely across biological membranes (Vermaas et al. 2019) making lignin oxidative polymerisation by LACs a potential main driving force controlling the metabolic gradient-dependent transport (Perkins et al. 2022). Here, we provide clear evidence for the fundamental mechanistic role of LACs in regulating lignin amount and composition differently between cell types and cell wall layers. In complement to LAC11, our study showed that LAC4 and LAC17, previously shown to be involved in vascular lignification (Berthet et al. 2011; Zhao et al. 2013), had non-redundant roles in the lignification of specific vascular cell types. Far from being the only actors, we also revealed novel and specific functions of LAC5, LAC10 and LAC12 in vascular lignification. In higher-order *lac* mutants, the loss of different LAC combinations resulted in specific and non-redundant biochemical changes on lignin chemistry affecting specific cell type and/or cell wall layer (figures 3–5). Table 1 summarises the different morphological and biochemical aspects observed in our higher *lac* mutants, which confirm the substrate-specific, layer-specific and cell type-specific activities of each of these different LACs. The drastic reduction in lignin contents in higher-order *lac* mutants, especially in fibre cell walls, demonstrated the implication of the five tested *LAC*s for vascular lignification (figure 4B). In contrast, the stable LAC activity in MX TEs in our higher-order *lac* mutant series suggests that the remaining *LAC11*, shown to be required for vascular lignification in the absence of *LAC4* and *17* (Zhao et al. 2013), suffices to ensure some lignification although both composition and function of TEs are impaired. TEs thus required specific lignins, differing between morphotypes, and accordingly LAC optimal pH and substrate ranges differed between PX and MX (figures 2 and S3). *Q* specifically affected the high pH-dependent activity present in MXs but absent in PXs when using the phenolic substrate pyrogallol (figure S3). Phenolic substrates have reduced redox potentials at higher pH, making them easier to oxidise (Rodgers et al. 2010). This likely explains the observed second activity peak for PYGL at pH 7 (figure S3), which is similar to the pH of the xylem sap in TEs (Blaschek and Pesquet 2021). This high pH combined with the high LAC activity maintained weeks after MX TEs have undergone cell death highlights the long-lasting capacity of sap conducting dead MX TEs to efficiently lignify *post-mortem* (Pesquet et al. 2013; Ménard et al. 2022). PRXs have also been shown to affect stem lignification (Blaschek and Pesquet 2021) and to have higher affinity for **S** than **G** monomers (Shigeto and Tsutsumi 2016). Their residual activity may explain the increased **S**/**G** ratios in the cell walls of higher-order *lac* mutants although our results suggest that LACs are predominantly responsible for the oxidation of **G** monomers. However, the almost complete absence of lignin from the IF SCWs suggests that PRXs are unable to fulfil the role of LAC for lignin polymerisation in IFs, thereby reinforcing the idea that PRXs and LACs have non-redundant functions during lignin formation (Zhao et al. 2013). Our results also suggest that LACs discriminate both different ring structures and aliphatic tails, as LAC4 favoured the incorporation of **G**_CHOH_ whereas LAC17 accumulated more **G**_CHO_ (figure 4). This non-redundant activity of LAC paralogs between cell types and cell wall layers (figure 2) is moreover supported by recent protein modelling results showing that LACs have distinct protein structures affecting the position of key catalysing amino-acids, active site binding pocket volume, shape and accessibility (Blaschek and Pesquet 2021). Previous localisation studies showed the ubiquitous localisation of LAC4 and LAC17 in the cell walls of fibres and TEs using fluorescent protein fusion (Schuetz et al. 2014; Hoffmann et al. 2020) and immunodetection (Berthet et al. 2011). LAC12 was shown to belong to the microtubule-linked cargo targeted to TE SCWs (Derbyshire et al. 2015). Our results clearly localised the activity of different LAC paralogs in distinct cell wall layers and cell types, such as LAC12 in the CC of IFs (figures 2, 5). *In vitro* LAC activity assays using heterologously expressed LACs have moreover yielded unexpected results, with either LACs showing no activity (Sato et al. 2001), activities that were at odds with the enzyme’s *in vivo* effect (He et al. 2019), or showing activity only after adding water soluble xylem extracts to the reaction buffer (Zhuo et al. 2022). Performing activity assays directly *in situ* avoids these effects by maintaining the native reaction environment of LACs and provides results that directly reflect the function of the enzyme *in planta*. As such, our results functionally demonstrate the role of differentially localised and tightly cell wall-bound LACs in specific cell wall layers (Chou et al. 2018) for controlling the spatial distribution of lignin. The specific depletion of lignin autofluorescence in the CCs of *Q-5, Q-10* and *Q* has a striking resemblance to the unlignified CCs in the eastern leatherwood (Mottiar et al. 2020). The atypical lignification pattern in that species was suggested to be responsible for the high flexibility of its wood (Mottiar et al. 2020), suggesting that evolutionary – or biotechnological – modulation of cell wall layer-specific LAC activity could adjust cell–cell adhesion and the biomechanics of whole tissues without compromising the structural integrity of the individual cells. In addition to the spatial restriction of lignification, specific cell types in vascular tissues maintain distinct lignin composition and amounts across species. TE enrichment in **G** residues for example is conserved among all vascular plant species (Pesquet et al. 2019). This cell type specific function of lignin has been recently demonstrated for the different TE morphotypes that require a tight regulation of **G**_CHO_ to **G**_CHOH_ ratio to maintain the balance between SCW stiffness and flexibility (Ménard et al. 2022). Here, we expand on the importance of distinct lignin chemistries by showing that specific LACs control lignin composition to ensure lignin function in structurally reinforcing TEs and regulating cell wall swelling in fibres (figures 6, 7). Altogether, our data provides the missing link between lignin monomer biosynthetic control and their polymerisation into distinct lignin polymers in specific cell types for plants to adapt to the numerous environmental and developmental stresses faced by their cell walls.

**Table 1.**
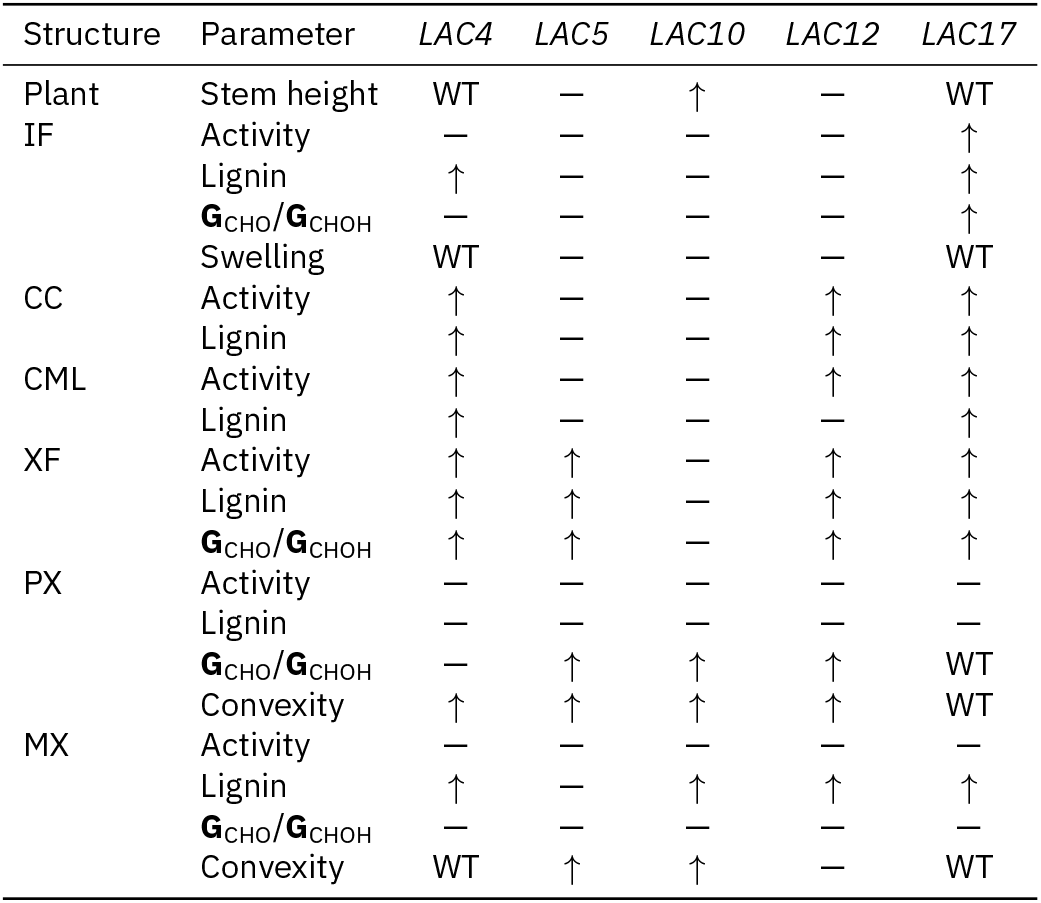
Effects of single active LACs in the *Q* background (*e.g.* the effect under *LAC4* describes the difference between *Q* and *Q-4*). –, no effect; ↑, partly restored; WT, restored to WT level.

| Structure | Parameter | *LAC4* | *LAC5* | *LAC10* | *LAC12* | *LAC17* |
| --- | --- | --- | --- | --- | --- | --- |
| Plant | Stem height | WT | — | ↑ | — | WT |
| IF | Activity | — | — | — | — | ↑ |
|  | Lignin | ↑ | — | — | — | ↑ |
| | $G_{CHO}/G_{CHOH}$ | — | — | — | — | ↑ |
|  | Swelling | WT | — | — | — | WT |
| CC | Activity | ↑ | — | — | ↑ | ↑ |
|  | Lignin | ↑ | — | — | ↑ | ↑ |
| CML | Activity | ↑ | — | — | ↑ | ↑ |
|  | Lignin | ↑ | — | — | — | ↑ |
| XF | Activity | ↑ | ↑ | — | ↑ | ↑ |
|  | Lignin | ↑ | ↑ | — | ↑ | ↑ |
| | $G_{CHO}/G_{CHOH}$ | ↑ | ↑ | — | ↑ | ↑ |
| PX | Activity | — | — | — | — | — |
|  | Lignin | — | — | — | — | — |
| | $G_{CHO}/G_{CHOH}$ | — | ↑ | ↑ | ↑ | WT |
|  | Convexity | ↑ | ↑ | ↑ | ↑ | WT |
| MX | Activity | — | — | — | — | — |
|  | Lignin | ↑ | — | ↑ | ↑ | ↑ |
| | $G_{CHO}/G_{CHOH}$ | — | — | — | — | — |
|  | Convexity | WT | ↑ | ↑ | — | WT |

## Author Contributions

EP conceived the study. EP, DM and LB designed the experiments. LB, DM and EM performed the experiments. LB and EP analysed the data. EP ensured financial support and scientific expertise. LB and EP wrote the article. All co-authors revised the manuscript.

## Acknowledgements

This work was supported by Gunnar Öquist fellowship from the Kempe foundation (to EP), Vetenskapsrådet (VR) research grants 2010-4620 and 2016-04727 (to EP), the Stiftelsen för Strategisk Forskning ValueTree (to EP), the Bolin Centre for Climate Research RA3, RA4 and RA5 “seed money” and “Engineering Mechanics for Climate Research” (to EP), and the Carl Trygger Foundation CTS 16:362/17:16/18:306/21:1201 (to EP). We also thank Bio4Energy (a strategic research environment appointed by the Swedish government), the UPSC Berzelii Centre for Forest Biotechnology, and the Departments of Materials and Environmental Chemistry (MMK), of Ecology, Environment and Plant Sciences (DEEP) and the Bolin Centre for Climate Research of Stockholm University (SU).

