## Supplementary material for "Different combinations of laccase paralogs non-redundantly control the lignin amount and composition of specific cell types and cell wall layers in *Arabidopsis*"

[1]Arrhenius laboratories, Department of Ecology, Environment and Plant Sciences (DEEP), Svante Arrhenius väg 20A, Stockholm University, 160 91 Stockholm, Sweden
[2]Umeå Plant Science Centre (UPSC), Department of Plant Physiology, Umeå University, 901 87 Umeå, Sweden
[3]Present address: Carlsberg Research Laboratory, J.C. Jacobsens Gade 4, DK-1799 Copenhagen V, Denmark
[4]Bolin Centre for Climate Research, Stockholm University, 106 91 Stockholm, Sweden

0000-0003-3943-1476 (LB), 0000-0002-6959-3284 (EP)

### Supplementary figures

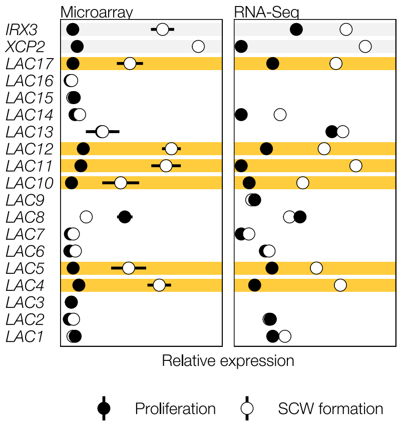


**Figure S1 | Additional information related to the choice of *LAC*s studied in this paper.** Co-expression of *LACCASE* genes with secondary cell wall formation in TEs. *XCP2* and *IRX3* are as marker genes expressed during SCW formation. Data is from Derbyshire et al., 2015 (doi: 10.1105/tpc.15.00314).

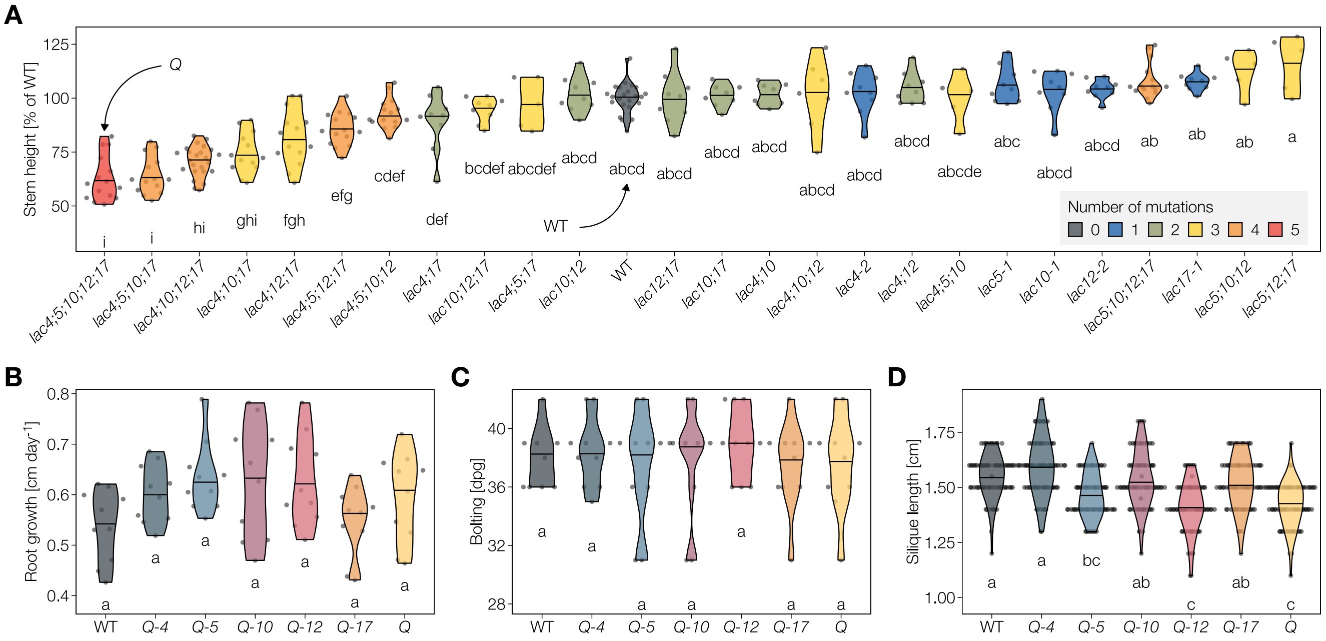


**Figure S2 | Additional information related to figure 1.** Phenotypical characterisation of higher-order laccase mutants. **A** Stem height at harvesting of the WT and *lac* single, double, triple, quadruple and quintuple mutants (colour coded), sorted by median plant height. $n$ = 5–24 individual plants per genotype. **B** Early root growth rate between 2 and 7 days past germination; $n$ = 10 individual plants per genotype from 2 plates per genotype. **C** The bolting time (defined as reaching a height > 5 cm) of higher-order *lac* mutants; $n$ = 10 individual plants per genotype from 2 independent growth instances. **D** Length of mature siliques from the main stem at time of harvest; 5 mature siliques were measured for each of $n$ = 15 individual plants per genotype from 3 independent growth instances. Different letters indicate statistically significant differences between genotypes according to a Tukey-HSD test (per panel; $\alpha$ = 0.05).

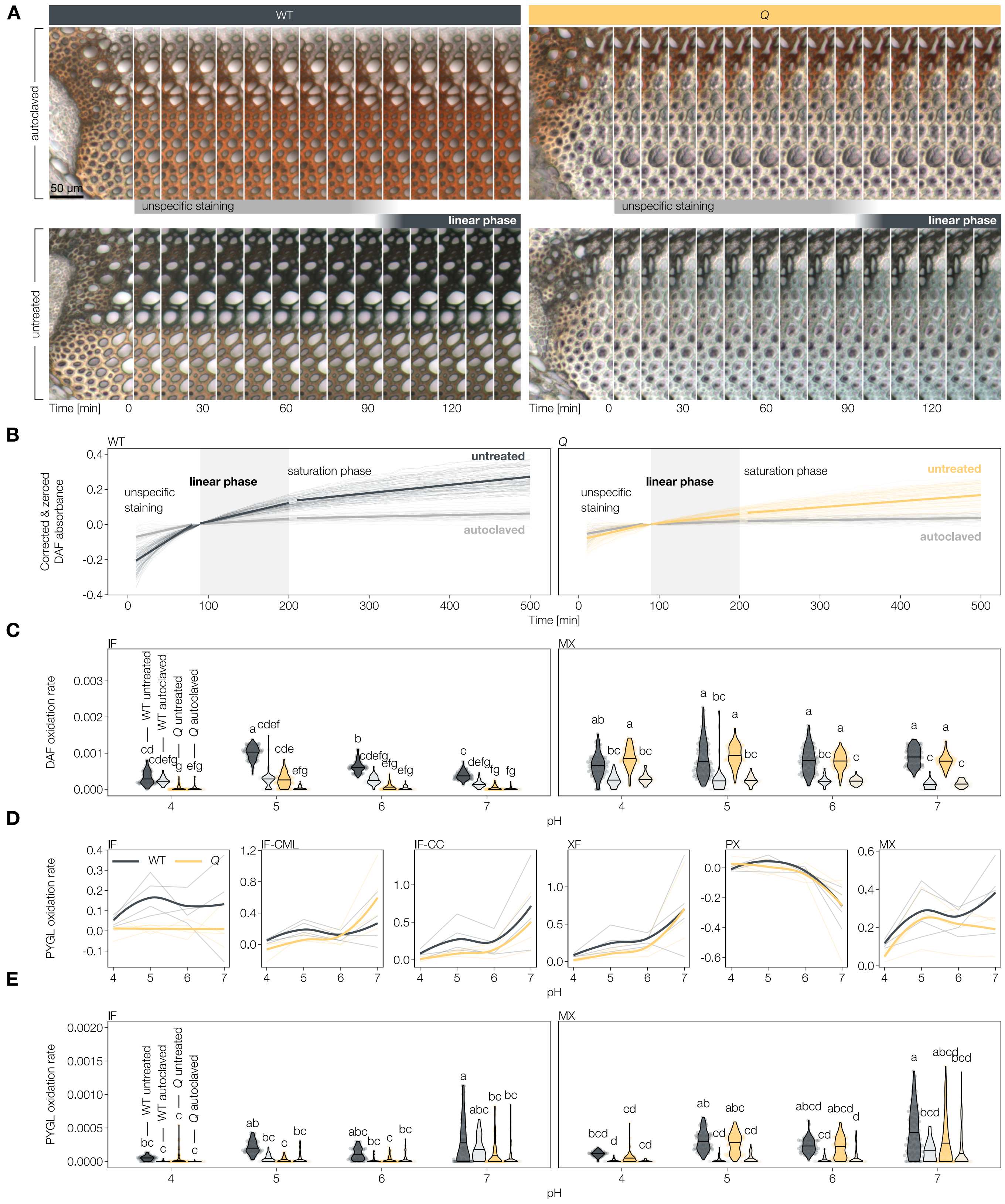


**Figure S3 | Additional information related to figure 2.** Time-course imaging of *in situ* LAC activity assays. **A** Real-time sequential image montage of autoclaved and untreated WT and *Q* sections incubated with DAF at pH 5. **B** Relative concentration of oxidised substrate in the cell wall of XFs measured during real-time imaging. Thin lines show 100 measured cell walls from 5 individual plants per genotype and condition. Thick lines show linear regressions of the initial phase of unspecific cell wall staining, the linear phase of substrate oxidation, and the saturation phase, in which activity levels off. **C** Relative DAF oxidation rates in the IF and MX of autoclaved and untreated WT and *Q* sections; n = 5 individual plants for untreated sections and n = 3 individual plants for autoclaved sections. **D** Optimal pH for activity towards the phenolic LAC substrate pyrogallol, showing an additional activity peak at pH 7 for multiple cell types. Activity in autoclaved sections is subtracted to show only LAC mediated oxidation. The activity rate drops into the negative when the apparent oxidation rate is lower than that of the unlignified phloem used for background correction. Thin lines represent 4 individual plants, thick lines are the overall average by local regression (LOESS). **E** Relative PYGL oxidation rates in the IF and MX of autoclaved and untreated WT and *Q* sections; n = 4 individual plants for untreated sections and n = 3 individual plant for autoclaved sections.
Different letters indicate statistically significant differences between genotypes according to a Tukey-HSD test (per panel; alpha = 0.05).

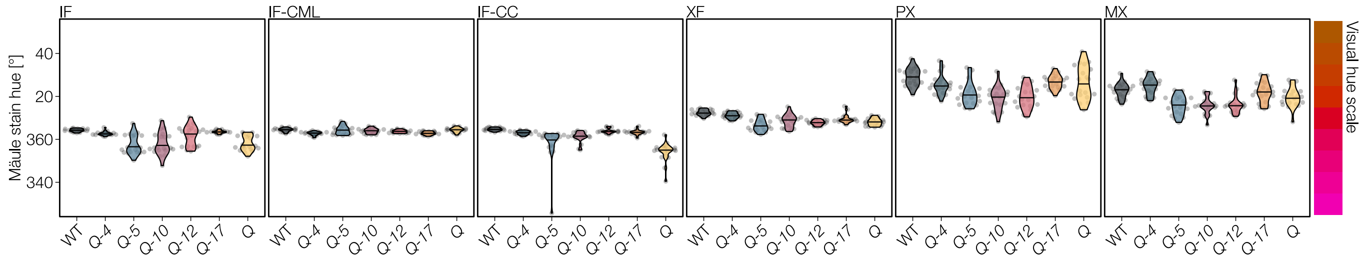


**Figure S4 | Additional information related to figure 3.** Hue of the Mäule stained cell walls of higher-order *lac* mutants. Twenty measurements per cell type and genotype from *n* = 1 stained section. Note that the quantitative analysis of the Mäule hue has never been validated and is an indication only.

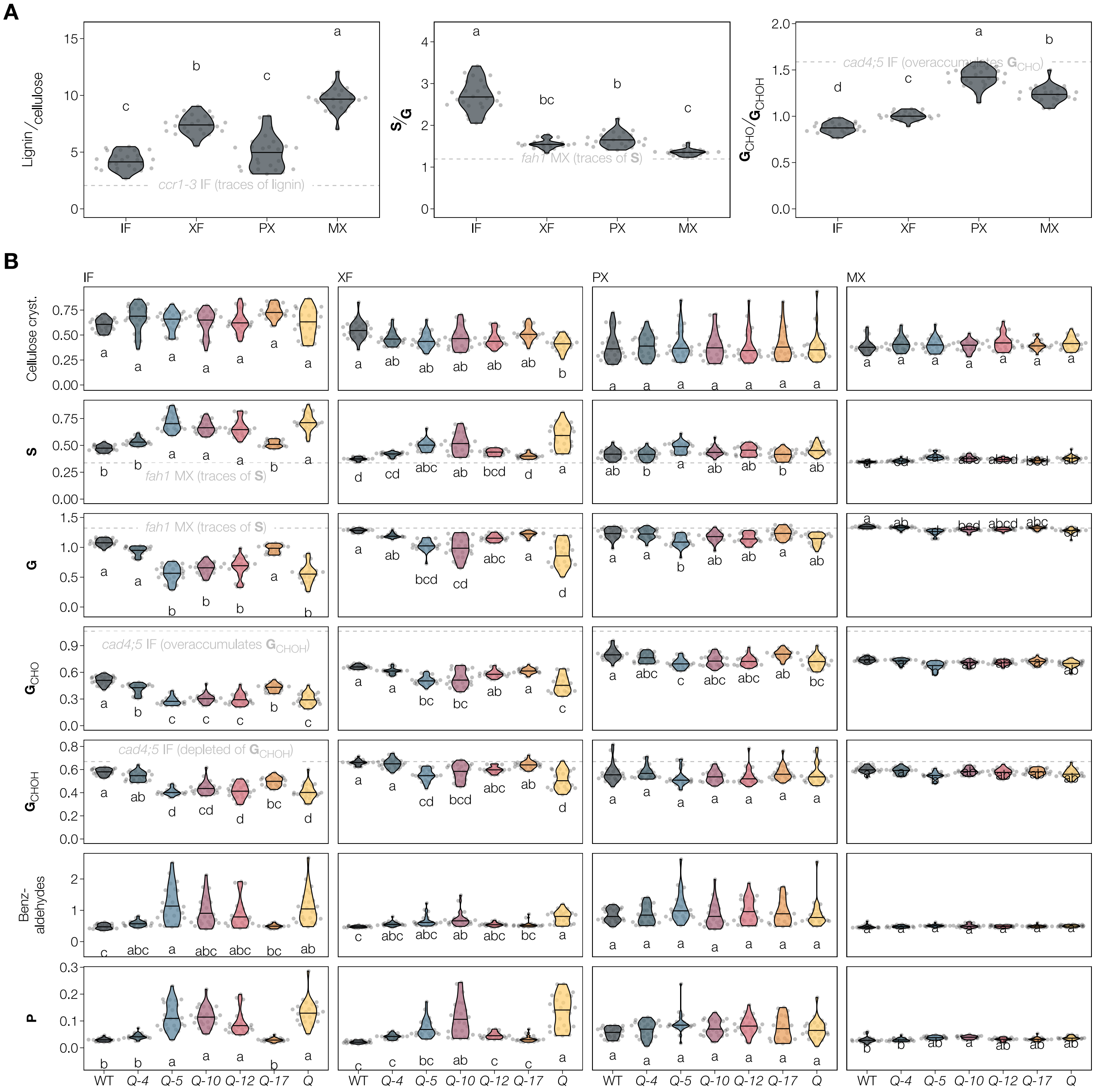


**Figure S5 | Additional information related to figure 4.** Lignin composition measured by Raman microspectroscopy. **A** Comparison of lignin quantity and quality between different cell types and morphotypes in WT plants. **B** Cell wall characteristics of WT and higher-order *lac* mutants. Different lignin constituents are expressed relative to total lignin amount. References from well characterised phenylpropanoid loss-of-function mutants are indicated by dashed grey lines to ease interpretation. Cellulose crystallinity is estimated as the ratio of the 378 and 1095 $cm^{-1}$ bands. Different letters indicate statistically significant differences between genotypes according to a Tukey-HSD test (per panel; $\alpha$ = 0.05); spectra of 5 individual cells were measured for each cell type in each of $n$ = 5 individual plants per genotype from 2 independent growth instances.

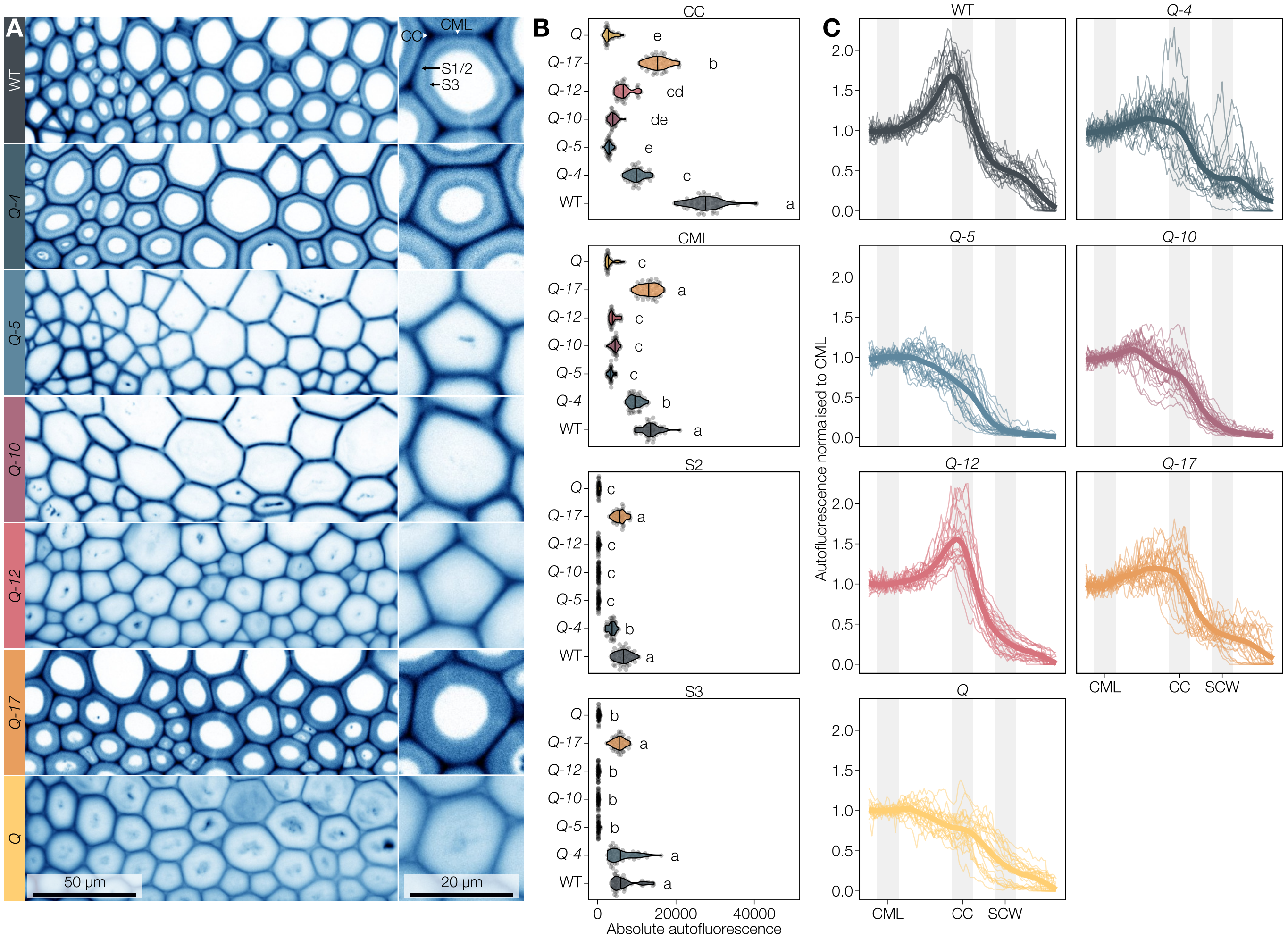


**Figure S6 | Additional information related to figure 5.** Cell wall layer-specific change in lignin autofluorescence. **A** Lignin autofluorescence in the IFs WT and mutant plants. Image contrast has been adjusted for each genotype for visibility. **B** Absolute lignin autofluorescence in CC, CML, S1/S2 and S3 layers of the IF in WT and higher-order *lac* mutants. The points represent 10 measurements per cell wall layer and genotype from each of $n$ = 5 individual plants per genotype from 2 independent growth instances. Different letters indicate statistically significant differences between genotypes according to a Tukey-HSD test (per panel; $\alpha$ = 0.05); **C** Autofluorescence intensity line profiles through CML, CC and SCW. Each line is normalised to the average of its CML. Thin lines are individual profiles from 5 independent plants, the thick line is an average by local regression (LOESS).

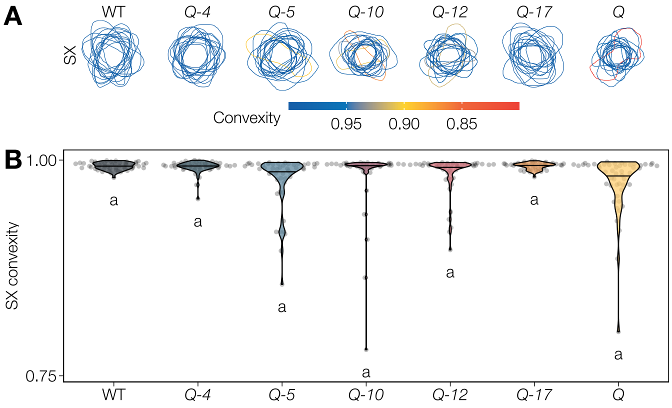


**Figure S7 | Additional information related to figure 6.** TE perimeters and their degree of inwards collapse in the SX of different higher-order *lac* mutants. Ten TEs in each of $n$ = 5 individual plants per genotype were measured. **A** The drawn outlines of the cell walls of fifteen randomly sampled TEs per genotype. **B** Degree of SX collapse in the different *lac* mutants and WT plants. Different letters indicate statistically significant differences between genotypes according to a Tukey-HSD test (per panel; $\alpha$ = 0.05).

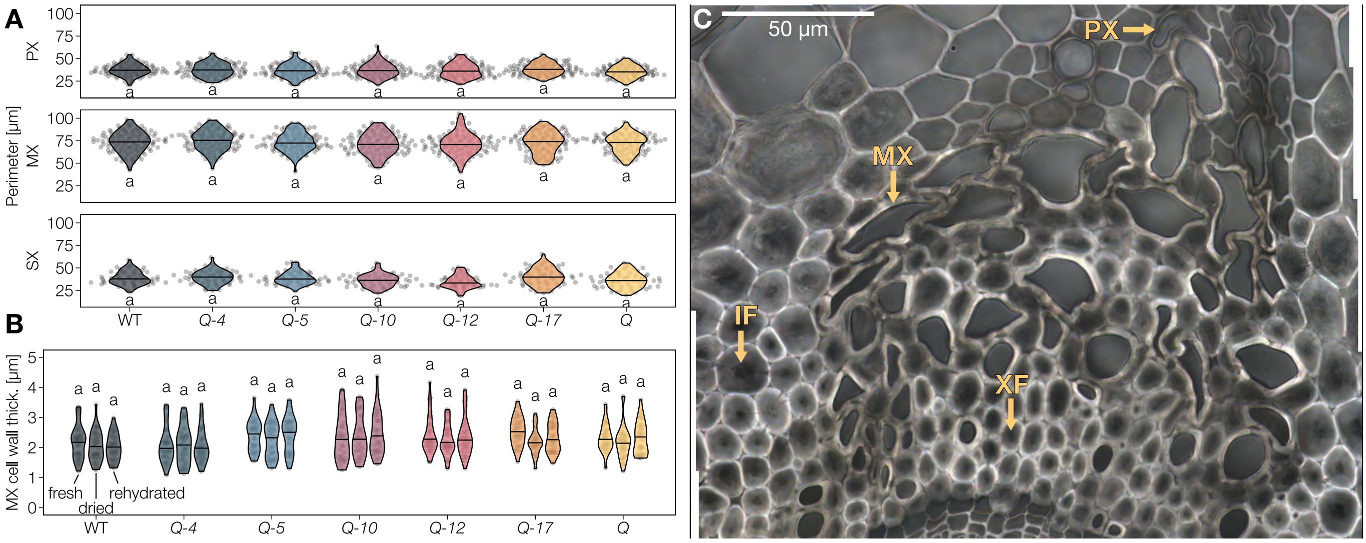


**Figure S8 | Additional information related to figure 7.** Structural integrity of cell walls. **A** Perimeter lengths of MX and PX TEs are undistinguishable in the WT and all tested mutants. Values are from the twenty TEs of each morphotype in each of $n$ = 5 individual plants per genotype shown in figure 6. **B** Thickness of cell walls is unaffected by drying and loss of *LAC* activity. Dots represent 10 TE cell walls from each of $n$ = 3 individual plants per genotype in the three states. Cell walls were measured lumen to lumen in TE–TE and TE–parenchyma cell walls, explaining the overall variation in cell wall thickness. Different letters indicate statistically significant differences between genotypes and states according to a Tukey-HSD test (per panel; $\alpha = 0.05$). **C** Differential interference contrast (DIC) image of a *Q* vascular bundle, showing the swollen cell walls in both IFs and XFs as well as inwardly collapsed TEs.

### Supplementary tables

**Table S1 |** Primers used for genotyping the *lac* mutant plants. Band sizes estimated from gel electrophoresis.

| Allele | Locus | Polymorphism | Insertion | WT allele primers (5'→3') | Mutant allele primers (5'→3') |
|---|---|---|---|---|---|
| *lac4-2* | AT2G38080 | GK-720G02-025278 | exon 3 | FW: TGGTAACTTTGGACGATCAGG<br>RV: AGTAATGAACAGTTGCGGTGG<br>band size: ~1 kb | FW: ATATTGACCATCATACTCATTGC[a]<br>RV: AGTAATGAACAGTTGCGGTGG<br>band size: ~1.1 kb |
| *lac5-1* | AT2G40370 | SALK_063466 | exon 5 | FW: ACTTCTCGTCTTTCCTCCTGC<br>RV: CTTGGAAGAGCAAATGAAACG<br>band size: ~1 kb | FW: ACTTCTCGTCTTTCCTCCTGC<br>RV: GCGTGGACCGCTTGCTGCAACT[b]<br>band size: ~800 bp |
| *lac10-1* | AT5G01190 | SALK_017722 | exon 5 | FW: CTATGGATCAATCAGAAGTCCG<br>RV: TCAATTCCAAGACATATCCGG<br>band size: ~1 kb | FW: CTATGGATCAATCAGAAGTCCG<br>RV: GCGTGGACCGCTTGCTGCAACT[b]<br>band size: ~750 bp |
| *lac11-1* | AT5G03260 | SALK_063746 | exon 5 | FW: ATTTCAATGTGACCGGACAACG<br>RV: TAAGTCTTGTCCCCGTTGATG<br>band size: ~1.2 kb | FW: GCGTGGACCGCTTGCTGCAACT[b]<br>RV: TAAGTCTTGTCCCCGTTGATG<br>band size: ~500 bp |
| *lac12-2* | AT5G05390 | SALK_125379 | exon 1 | FW: AGGGAAAGGAAAAGAGGAACC<br>RV: TTTCTGCCAACATTTTTGAGG<br>band size: ~1.1 kb | FW: GCGTGGACCGCTTGCTGCAACT[b]<br>RV: TTTCTGCCAACATTTTTGAGG<br>band size: ~500 bp |
| *lac17-1* | AT5G60020 | SALK_016748 | promoter | FW: TCGAAGAGGGTCAAAGAGTTT<br>RV: TCTTAGCCATGAAATGTGAGC<br>band size: ~900 bp | FW: GCGTGGACCGCTTGCTGCAACT[b]<br>RV: TCTTAGCCATGAAATGTGAGC<br>band size: ~900 bp |

[a] Gabi-Kat LB primer
[b] SALK LB primer

**Table S2 |** Synthetic substrates used for activity assays in sections.

| Name | Abbreviation | Supplier | Identifier | Stock concentration | Solvent | Working conc. |
|---|---|---|---|---|---|---|
| 2,7-Diaminofluorene | DAF | Sigma-Aldrich | D17106 | 7 mM | 24 mM HCl in water | 70 μM |
| 3,3'-Diaminobenzidine | DAB | Sigma-Aldrich | D8001 | 5 mM | 12 mM HCl in water | 500 μM |
| Pyrogallol | PYGL | Sigma-Aldrich | P0381 | 50 mM | Water | 5 mM |
